# Ring Array Illumination Microscopy: Combination of Super-Resolution with Large Field of View Imaging and long Working Distances

**DOI:** 10.1101/2023.09.05.555896

**Authors:** Johann von Hase, Udo Birk, Bruno M. Humbel, Xiaomin Liu, Antonio V. Failla, Christoph Cremer

## Abstract

Here we present a novel fluorescence microscopy concept which enables a direct integration of Super-Resolution Microscopy (SRM) approaches (SIM/Nanosizing, STED, SMLM, MINFLUX, SIMFLUX) into microscopy systems with working distances (WD) up to the multicentimeter range while still allowing nanometer scale resolution at selected sites. This becomes possible by a “synthetic aperture” illumination mode with multiple, constructively interfering excitation beams positioned in a “Ring-Array” arrangement around a beam free interior zone containing instrumentation involved in complementary imaging modes. The feasibility of such a direct correlative microscopy method is validated by extensive numerical simulations; on the basis of these calculations, experimental implementation options are discussed. Such “Ring Array” illumination modes may be useful for various correlative microscopy methods, such as a direct combination of correlative light and electron microscopy in the same device (dCLEM); or a direct combination of low NA/large field-of-view widefield microscopy and super-resolution of selected sites in the same device (direct Correlative Opical Microscopy/dCOLM). Ring-Array supported correlative microscopy modes will open novel imaging perspectives in a variety of disciplines, from material sciences to biomedical applications.

## Introduction

In many applications of microscopy in the biosciences as well as in the material sciences, it is highly desirable to combine a large field of view (FOV)/long working distance screening provided at low optical resolution, with the enhanced resolution imaging (ERI) of selected sites. This can be obtained by a variety of means. Fluorescence based super-resolution microscopy (SRM) methods, for example, presently achieve best optical resolution values down to the single molecule level; transmission and scanning electron microscopy, focused ionized particle beam microscopy, X-ray microscopy, or X-ray tomography (hereafter referred to as “ultrastructural microscopy”), in some cases can reveal fine structural details even down to atomic resolution.

In conventional light microscopy, the correlative combination of low resolution and high resolution imaging modes has been in use already since the 1850s by the invention of the objective lens revolver. Using current systems of SRM, however, the change of the objective lens produces mechanical and thermical modifications at the nanoscale which may require time consuming recalibration of the SRM system (Schaufler, 2017).

In other applications, the ERI mode of selected sites identifed by low resolution/large WD microscopy typically is performed by a transfer of the specimen to a physically separated microscopy system. In alternative methods of directly Correlated Electron Light Micorscopy (dCLEM), low resolution light microscopy is used to identify the target (region of interest/ROI), while electron microscopy (or other ultrastructural microscopy techniques such as X-ray tomography) provide the high-resolution image data and the structural context. The better the optical resolution (object plane coordinates: x, y; coordinates along the optical axis: z) of the light microscope system used, the more useful the dCLEM technique becomes. In addition, light microscopy and electron microscopy/ultrastructure microscopy are complementary techniques for structure elucidation due to different contrast mechanisms; the better the optical resolution of light microscopy, the more informative and useful the combination of these two optical approaches.

To overcome the experimental difficulties encountered in present approaches of correlated light optical microscopy (CLOM) and correlated light electron microscopy (CLEM), it should be highly useful to integrate super-resolution methods directly into the low resolution light optical imaging system (dCLOM) or the ultrastructural system (dCLEM). In the following, we describe a concept which allows such a direct implementation by the realization of a “Ring-Array Microscopy” mode.

### Super-resolution light microscopy

Due to novel developments in optical technology and photophysics (Hell 2009; Cremer 2012; Cremer & Masters 2013; Ehrenberg 2014; Sydor et al. 2015; Lelek et al., 2021), it has become possible to radically overcome the classical diffraction limit for high numerical aperture (NA) objective lenses (approximately 200 nm in the object plane, 600 nm along the optical axis) of conventional far-field fluorescence microscopy (Abbe 1873; Rayleigh 1896). Examples of these methods of enhanced resolution of far-field fluorescence microscopy by “super-resolution” microscopy/ SRM (Toraldo di Francia, 1955) include 4Pi microscopy (Cremer & Cremer 1978; Hell et al. 1994a; Hänninen et al. 1995); localization microscopy using quantum dots (Lidke et al. 2005) or nanographenes (Xiaomin et al. 2020); photoactivated proteins (Betzig et al. 1995, 2006; Hess et al. 2006; Lemmer et al. 2008) or standard fluorophores (Cremer et al 1996, 1999; Esa et al. 2000; Rust et al. 2006; Reymann et al. 2008; Heilemann et al. 2008); stimulated emission depletion (STED) microscopy (Hell & Wichmann 1994; Hell et al. 1999; Hell 2009); or structured illumination microscopy (Heintzmann & Cremer 1999; Gustafsson 2000; Albrecht et al. 2002; Baddeley et al. 2007). Different modifications of localization microscopy (here referred to as SMLM) have been called by different denominations (Masters & Cremer, 2013), e.g., pointillism, PALM, FPALM, SPDM, STORM, dSTORM, etc.. In the context of Ring-Array microscopy, all of these “super-resolution microscopy” (SRM) methods can be applied. In the aforementioned SRM approaches, both the optical resolution (smallest detectable distance between two adjacent point sources according to relations (1) and (2)) and the structural resolution (smallest structural detail determined, for example, based on the density of the resolved point sources; Birk et al., 2017) may be very substantially enhanced.

At the current state of the art, these SRM techniques, using high numerical aperture (NA) objective lenses, currently allow light-optical resolution of biostructures down to about 5 nm (Galbraith and Galbraith 2011; Cremer 2012; Cremer et al. 2011), corresponding to about 1/100 of the wavelength λ_exc_ used for fluorescence excitation. Recently, a laser scanning microscopy-based SRM technique called MINFLUX (Balzarotti et al. 2017; Gwosch et al. 2020) based on a combination of donut-like illumination and enhancement of localization has been described; in this way, a which light-optical resolution down to the 1 nm range may be realized. According to the state of the art, this latter method also requires high numerical aperture (NA) objective lenses.

In the following, SRM techniques are understood to comprise all lightoptical approaches enabling enhanced lateral optical resolution beyond that provided by the relation

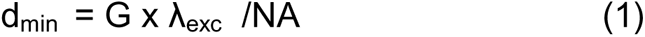

Here, λ_exc_ is the wavelength in vacuum used for fluorescence excitation, NA is the numerical aperture of the objective lens applied; “lateral” is the optical resolution in the object plane; NA = n sinα, where n is the refractive index of the material and α is half the aperture angle between a point of the object and the front lens of the objective lens used. Accordingly, in the Ring-Array illumination method described here, α is also understood to be half of the aperture angle between a point of the object and the Ring-Array arrangement (see Fig. 1).

**Fig. 1:**
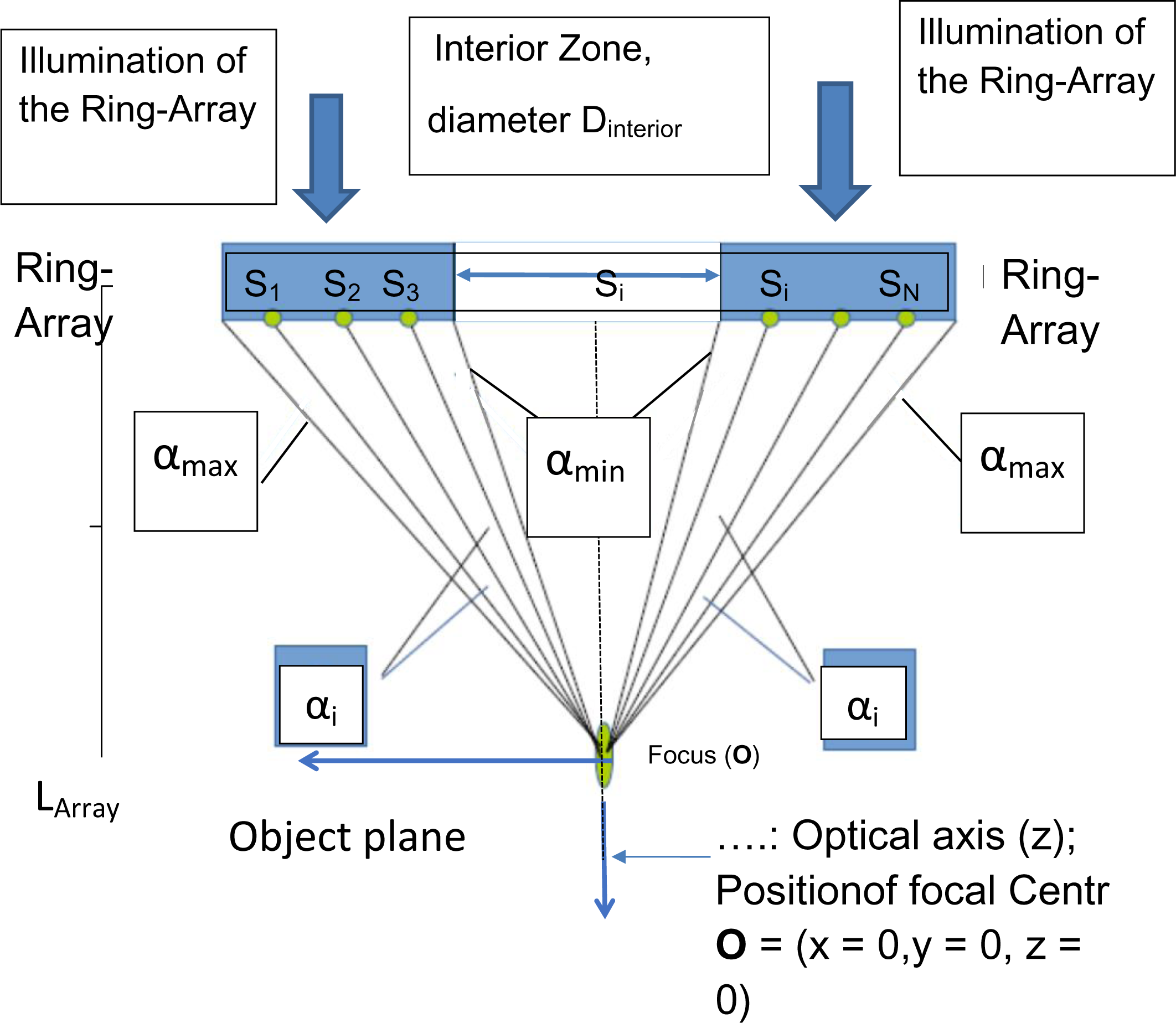
General scheme of Ring-Array Illumination. Individual sources S_1_, S_2_,… S_i_,.. S_N_ of coherent, e.g. collimated light waves are positioned at positions X_i_ = (x_i_, y_i_) in a flat illumination ring zone (hereinafter referred to as ring-array); for each S_i_ with individually defined phase, power and polarization relationships, coherent light is emitted from the S_i_ at angles α_i_ in defined directions (beam direction indicated by the direction of the wave vector). The illumination sources can be arranged in different ways in the illumination Ring-Array, see e.g. Fig. 4.

Far-field microscopy methods that result in enhanced resolution compared to (1) (i.e., d_min_ < G x λ_exc_/NA) are referred to as “nanoscopy” or “super-resolution microscopy” (SRM) methods (Cremer 2012; Cremer & Masters, 2013). Here, the term “objective lens ” refers to an optical device consisting of elements transparent to light (visible to near-ultraviolet spectral range), typically containing one or more lenses made of glass or other transparent materials. For the relationship of the lateral half-width (Full-Width-at-Half Maximum/FWHM_latera_l) of the corresponding lateral point spread function (PSF) with the lateral optical resolution according to relation (1), the factor G = 0.61 or 0.51 is usually assumed; depending on the application, slightly different factors G are also possible. “PSF” here refers to the normalized intensity distribution of a “point-like” (diameter << wavelength λ_exc_) fluorescent object. Since these minor differences are not relevant for the general concept of Ring-Array microscopy, example calculations are based on relationship (1) with factor G = 0.61 or FWHM_lateral_ = 0.51.

As an estimate of the conventional optical resolution of far-field fluorescence microscopy using a single objective lens along the optical axis of the microscope system, the estimates

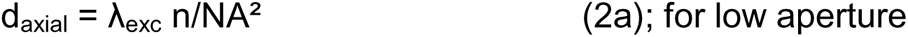

or

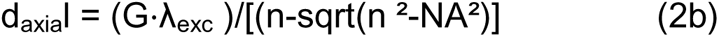

are used, where G ≈ 0.88 is assumed, and

λ_exc_, n, NA have the same meaning as in relation (1).

For other estimates of d_axial_ and FWHMaxial (FWHM along the optical axis of the microscope system), a factor slightly different from 2 is obtained in each case (between 1 and 2); however, since these small differences are not relevant to the Ring-Array concept discussed here, for simplicity of argument example calculations for d_axia_l and FWHM_axial_ are based on relation (2a).

Light-optical microscopy techniques with enhanced axial resolution with respect to relation (2) are also subsumed under the term “super-resolution”. This is the case, for example, for confocal laser scanning 4Pi fluorescence microscopy based on two opposing objective lenses with high NA (Hell & Stelzer 1992; Hell et al. 1994a,b), which, compared with the conventional estimate (e.g., λ_exc_ = 488 nm; NA = 1.4; n = 1.515) of d_axial_ = 488 nm x 1,515 /1.4^2^ = 380 nm (according to Eq. 2a), or d_axial_ = 0.88 x 488 nm/[(1.515 - sqrt(1.5152^2^ - 1.4^2^)] = 460 nm (according to Eq. 2b), respectively, result in a much improved axial resolution (FWHM_axial_) of about 80 - 100 nm.

In the context of the Ring-Array Illumination concept it is of minor relevance to which extent relations 2a) to 2c) may be more accurate with respect to typical experimentally observed axial FWHMs: Both relations 2(a,b) indicate a very substantial dependence on the Numerical Aperture. For example, assuming NA = 0.1, n = 1, G = 0.88 and λ_exc_ = 488 nm, relation 2a) gives d_axial_ =48.8 µm; for 2b), d_axial_ = 85.7 µm is obtained.

Furthermore, in all numerical exemples presented here, unless explicitly stated otherwise, λ_exc_ = 488 nm is used for the excitation wavelength and n = 1 (vacuum) for the refractive index between the object and the fluorescence detection system, or n = 1.31 for the refractive index of a sample embedded in ice or water. Example values for other excitation wavelengths and numerical apertures are easily obtained from relations (1) and (2) (Born & Wolf 1980; Feynman 2006).

As a measure of the three-dimensional optical resolution (”volume resolution”), the following relationship applies (Stelzer and Lindek 1994)

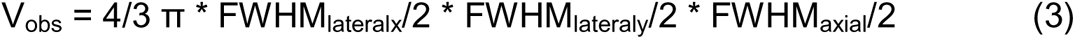

where

V_obs_ is called the observation volume,

With FWHM_lateralx_ the half-width of the PSF in the coordinate direction (x) of the object plane (x,y), (For the definition of the coordinate axes or their designations see Fig. 1);

FWHM_lateraly_ the half-width of the PSF in the coordinate direction of the object plane (x,y) orthogonal to (x), and

FWHM_axial_ represents the half-width of the PSF along the optical axis (z) of the optical system used for fluorescence detection.

### Correlative light and electron microscopy

Correlative light and electron microscopy (CLEM) is well established for sample preparation at room temperature (for a review, see Loussert, Fonta, and Humbel, 2015); but for cryo-electron microscopy, some challenges still need to be overcome. As an example, the application according to the state of the art to the analysis of biological samples is described below. Currently, in most cases, the samples of interest (typically cells) are mounted or grown on electron microscopic meshes and frozen. These, now frozen cells are first examined with a cryo-fluorescence microscope to find the structure of interest in the cell, such as an organelle. A focused ion beam (FIB) scanning electron microscope (SEM) is used to cut out a lamella in this area to make the sample thin enough for electron tomography (Sartori et al., 2005; Leis et al., 2006; Sartori et al., 2007; Schaffer et al., 2007; Rigort et al., 2012; Schaffer et al., 2015; Mahamid et al., 2016; Schaffer et al., 2017; Schorb et al., 2017). The workflow includes cryofixation, transfer of the sample to a cryo-light microscope, imaging with photons, transfer to a dedicated cryo stage of a FIB SEM, relocalization of the region of interest, preparation of the lamella and transfer to a cryo-transmission electron microscope, and imaging with electrons.

There are three transmission steps, and each step carries the risk of ice contamination, with the last step being the most vulnerable and important. Furthermore, (see above) currently the optical resolution in the object plane (x, y) of a conventional optical microscope is at best about 200 nm, while the z-resolution (along the optical axis) of the optical microscope used is at best only in the range of 500 nm; i.e., in the range of the maximum thickness of a sample that can be analyzed by transmission electron microscopy (TEM). It should be noted that these values apply only to high-resolution oil immersion objectives, which cannot be used because of the vacuum conditions in the electron microscope and the sample setup. For this reason, the state of the art makes it difficult to select the region of interest to be studied in FIB-SEM mode.

The use of a dedicated light microscope (use of oil objective lenses with high numerical aperture, e.g. NA = 1.4) outside the FIB-SEM leads to better x,y and z resolution at room temperature. Cryosamples, on the other hand, cannot be imaged with oil immersion objectives, which means that the numerical aperture (NA) cannot be larger than 1.

According to the state of the art, current FIB SEM techniques, as described above, can require long processing times to prepare the region of interest. In addition, for thick samples, the region of interest can be easily missed in the z-dimension, due to the poor z-resolution of the light optics.

Another direct imaging technology is cryo-FIB tomography (Schertel et al, 2013). After the region of interest is identified, a thin layer of it, a few nanometers thick, is etched away with the ion beam; then the newly created surface is imaged with the electron beam. This process is repeated until the entire region is imaged as a 3D volume. A typical volume is 60 x 40 x 20 μm^3^. Again, finding the area of interest suffers from the poor resolution of optical microscopy and the inaccurate overlay of the 3D map it creates. Direct control of the ion beam with an integrated high resolution optical microscope would greatly improve precision.

In a CLEM procedure where the light optical analysis is performed outside the FIB-SEM system (or other ultrastructure microscope), generally any microscopy setup can be used.

An alternative approach to perform CLEM with improved optical resolution is to use structured illumination light microscopy (SIM) according to the state of the art (Gustafsson 1999; Gustafsson et al. 2000; Heintzmann & Cremer 1999). In conjunction with a high numerical aperture (NA) objective lens (oil immersion), SIM allows a (x, y) resolution of about 100 nm and a z resolution of about 300 nm (for details see Suppl. Material). Using lower NA objective lenses, the SIM resolution is diminished but still provides superior contrast (Schneckenburger et al. 2020).

For a given numerical aperture (NA), SIM provides optical resolution enhanced by a factor of two; e.g. for NA = 0.2 (λ_exc_ = 488 nm, n = 1), this gives a theoretical lateral optical resolution of d_SIMmin_ = 0.6 µm laterally and d_SIMaxial_ = 6.1 µm; for NA = 0.1, the lateral optical resolution achievable with SIM is d_SIMmin_ (NA = 0.1) = [0.51λ_exc_/(NA]/2 = 1.2 µm; for the axial resolution, under these conditions (n = 1), d_SIMaxial_ = [λ_exc_ /NA²]/2 = 24 µm is obtained; for NA = 0.06 (working distance in the range suitable for integrated cryo-CLEM), a lateral resolution d_SIMmin_ (NA = 0.06, λ_exc_ = 488 nm) = 1.2 µm and an axial resolution d_SIMaxial_ (NA = 0.06) = 68 µm is estimated.

“Proof-of-principle” experiments with retina cells using a low NA SIM microscope with a distance from the object plane to the nearest optical element of the SIM device of about 4.5 cm (Best 2014; Schock et al., 2022) resulted in an optical lateral resolution of about 2 μm, in agreement with the theoretical estimate. The resolution and observation volume obtained was greatly enhanced compared to microscopy at the same distance but without SIM, but still far from the range desired for integrated correlative microscopy in a FIB SEM.

In many cases of correlative microscopy it may be sufficient to obtain more infromation about the size of an optically isolated small fluorescent object. Using axially structured illumination and high NA objective lenses, a size resolution down to few tens of nm has been obtained (Failla et al., 2002; Mathee et al. 2006).

In addition to SIM, other SRM approaches with improved resolution such as STED, single molecule microscopy (SMLM), MINFLUX, or other high-resolution imaging techniques (e.g., SIMFLUX) might be used. For example, according to the state of the art (Reymann et al., 2008; Lemmer et al., 2008, 2009, 2012; Cremer et al., 2017), relatively high illumination intensities (in the range of 1 - 10 kW/cm^2^) are typically used for optical resolution of single closely adjacent molecules. This is usually achieved by focusing laser radiation of wavelength λ_exc_ at powers in the range of a few hundred mW to object plane areas with diameters in the range of a few tens of µm.

In an integrated CLEM approach (dCLEM), where the light microscopic analysis of the sample is performed directly in the FIB-SEM, only objective lenses with large working distances (corresponding to a low Numerical Aperture) can be used due to geometric limitations. Similar limitations must also be considered when combining FIB-SEM directly with other ultrastructural microscopy methods or other imaging techniques.

### Correlated Light and Electron Microscopy (in the following called correlative microscopy)

For example, Gorelick et al. (2019) described an integrated cryocorrelative light and FIB-SEM microscope that allows direct and rapid identification of cell regions (ROIs) using fluorescence microscopy. In this case, an excitation and emission element was directly integrated into the FIB-SEM along with an objective lens for fluorescence imaging. However, due to the geometric constraints required by the instrumentation for FIB-SEM (compare Figure 2 of Gorelick et al. 2019), only a low NA objective lens was used, in this case NA = 0.06. According to relations (1) and (2), such an objective lens allows a lateral resolution (object plane, xy) of 4.2 µm and an axial resolution (along the optical axis (z) of the objective) of 135 µm (assumed excitation wavelength λ_exc_ = 488 nm; n = 1).

**Fig. 2:**
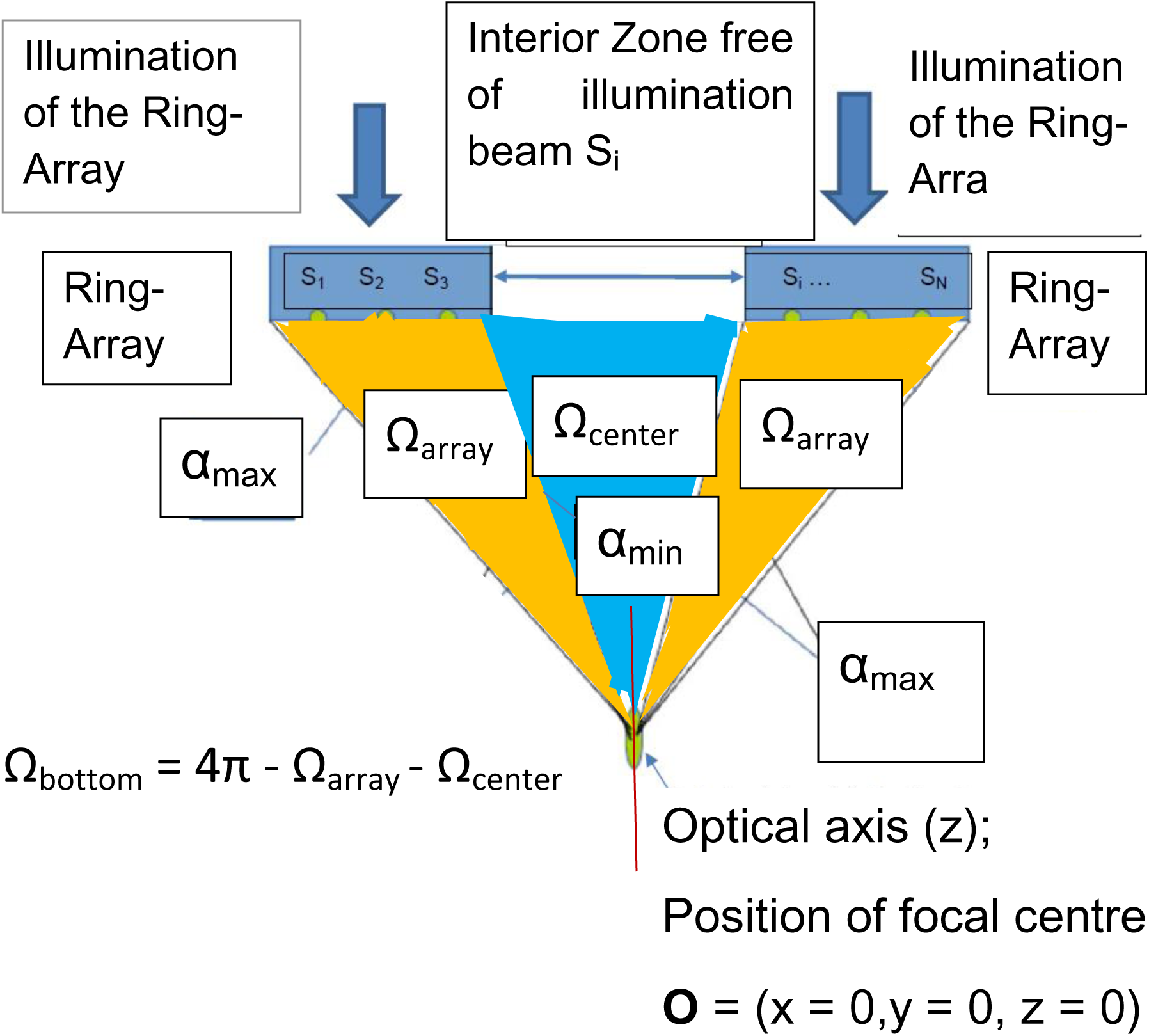
Identification of solid angles Ω_array_ and Ω_center_. Fig. 2 again shows the general arrangement of Fig. 1, but this time with special indication of the areas for the solid angles Ω_array_ (marked yellow) and Ω_center_ (marked light blue). The area filled by the solid angle Ω_bottom_ results from Ω_bottom_ = 4π - Ω_array_ - Ω_center_. To illuminate the Ring Array arrangement to generate the coherent light emitted by the sources S_i_ waves, lasers or other methods of generating sufficiently coherent radiation can be used.

For technical reasons, the numerical aperture (NA) of objective lenses with very large working distances is limited to low values: For objective lenses with high NA, geometrical optics requires that the radius of the optical lens (in practice, the front lens) be of the same order of magnitude as the working distance (Wilson and Sheppard 1984). Very large lenses are currently not only difficult (and expensive) to manufacture with the required imaging quality, but also difficult to accommodate and mount (aside from the severe geometric constraints within an FIB SEM). Second, the application of immersion embedding for large working distances is not straightforward: this would require the entire space between the front lens of the objective and the sample (ROI) to be filled by the immersion fluid, which is generally contrary to the application concept of large working distances. In the case of FIB-SEM, the sample must be introduced into a high vacuum chamber at very low temperature. Therefore, in most cases the use of immersion embedding not possible.

In principle, the undesired effects of the low resolution PSF necessarily obtained in low NA objective lenses might be overcome by using super-resolution approaches, such as for example Stimulated Emission Depletion (STED) microscopy:

Basically, the lateral optical resolution d_min_ in STED microscopy according to the state of the art is given by

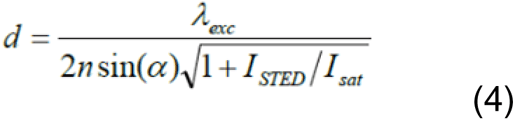

where λ_exc_ is the fluorescence excitation wavelength, NA = *n* sin (*α*) is the numerical aperture of the objective used, I_STED_ is the intensity of the donut-focused STED beam, and I_SAT_ is the saturation intensity of the fluorophore used for STED imaging (Hofmann et al. 2005). This relationship states that, in principle, it should be possible to achieve “any good” STED resolution at any numerical aperture (i.e., also at any working distance) by suitably increasing I_STED_; however, according to relationship (4), assuming the same wavelength and STED resolution for high resolution (e.g., d_minSTED_ = 50 nm), the required STED beam intensity scales inversely proportional to NA^2^, according to I_STED_ =I_SAT_ λ_exc_^2^/(4d_min_^2^ NA^2^) - 1. This means that with an objective lens with a sufficiently large working distance WD (e.g., for NA = 0.2), an about 50 times higher STED beam intensity I_STED_ (factor 1.4/0.2)^2^ would be required to achieve the same lateral super-resolution (e.g. 50 nm) as with NA = 1.4. For an objective lens with a numerical aperture of NA = 0.06 and a sufficiently large working distance for integrated correlative light and FIB SEM microscopy (Gorelick et al. 2019, see above), even an about 500 times higher STED beam intensity would be required to achieve the same super-resolution as with NA = 1.4.

Currently, typical I_STED_ intensities for high NA objectives are in the tens of MW/cm^2^ (Bordenave et al. 2016); this would result in I_STED_ intensities on the order of 5 GW/cm^2^ for high resolution low NA STED microscopy (e.g., NA = 0.06) suitable for integrated correlative microscopy. In many applications, such high values are unacceptable; bleaching and phototoxicity already result in many STED-Microscopy applications with high NA to adverse effects (Li and Betzig 2016); in the case of FIB-SEM, this likely would also lead to unacceptable heating of the sample.

Alternatively, it remains highly desirable to develop high-resolution (SRM) techniques for direct integration into correlated light and FIB SEM devices for sufficiently large working distances (up to the multicentimeter range) with much lower illumination intensities.

To summarize: What is needed to solve the above stated problems of great practical importance for correlative FIB-SEM applications as well as other applications with large working distance of the light-optical elements is a high-resolution light microscopy technique that enables the following:

a. the use of large working distances (WDs); for geometrical reasons, working distances (WDs) of several cm are required in a typical FIB SEM; large working distances are also desirable for screening large fields of view by low NA microscopy.
b. the identification of the target (object structure of interest, ROI) and its position within a transparent sample with an accuracy as close as possible to the section thickness of the FIB-SEM (e.g., 5 nm or 10 nm, or a few tens of nm), but in any case with an accuracy that substantially limits the number of sections required for further containment compared to the prior art (e.g., 50 sections of 10 nm each instead of 5,000 sections).

Similar arguments suppport the realization of enhanced resolution analysis of sites selected by low NA optical screening: Using low NA optics, a large field of view (e.g. 1 cm^2^) may be screened in a few seconds, selecting a few specific sites for SRM analysis; to screen e.g. 1 cm^2^ (= 10^14^ nm^2^) by STED at a desired resolution of 10 nm, would require scanning with a pixel size of 5 nm; assuming a dwell time of 1 µs/pixel, the scanning would still take ca. 50 days.

A basic concept for achieving superresolution at “arbitrarily” large working distances based on holographic laser scanning 4π-microscopy without the use of objective lenses was presented as early as the 1970s (C. Cremer and T. Cremer 1972, 1978); in this early concept of enhanced synthetic aperture imaging, one or more “point holograms” were proposed to achieve an observation volume (focal volume) much smaller than achievable with a conventional objective lens microscope of high NA; imaging was to be performed by “point-by-point” scanning of the object of the object through the focus thus created.

Instead of holograms, distributed aperture illumination arrays can also be applied (Birk et al. 2017): here, a limited number of individual collimated coherent beams are used, where intensity, direction, and phase are individually controlled compared to the very large number of diffracted beams (e.g., hundreds of thousands) generated by the hologram structures with coupled intensities, directions, and phases. This has the great advantage that the light sources can in principle be positioned at any distance from the scanning focus area (”spot”) where constructive interference occurs; by changing the direction of the collimated coherent beams accordingly, the scanning area (”spot”) can be moved to any position in 3D. If necessary, the phases and other parameters of the collimated beams must also be readjusted for the new position. However, these approaches did not take into account that in many applications, e.g., correlated FIB-SEM or numerous other microscopy techniques with the requirement of large working distances, a considerable solid angle (> 2π) around the optical axis (see Fig.1 - 3) cannot be used for correlative microscopic imaging. This considerable disadvantage of current techniques is eliminated by the Ring-Array illumination approach presented here.

**Fig. 3:**
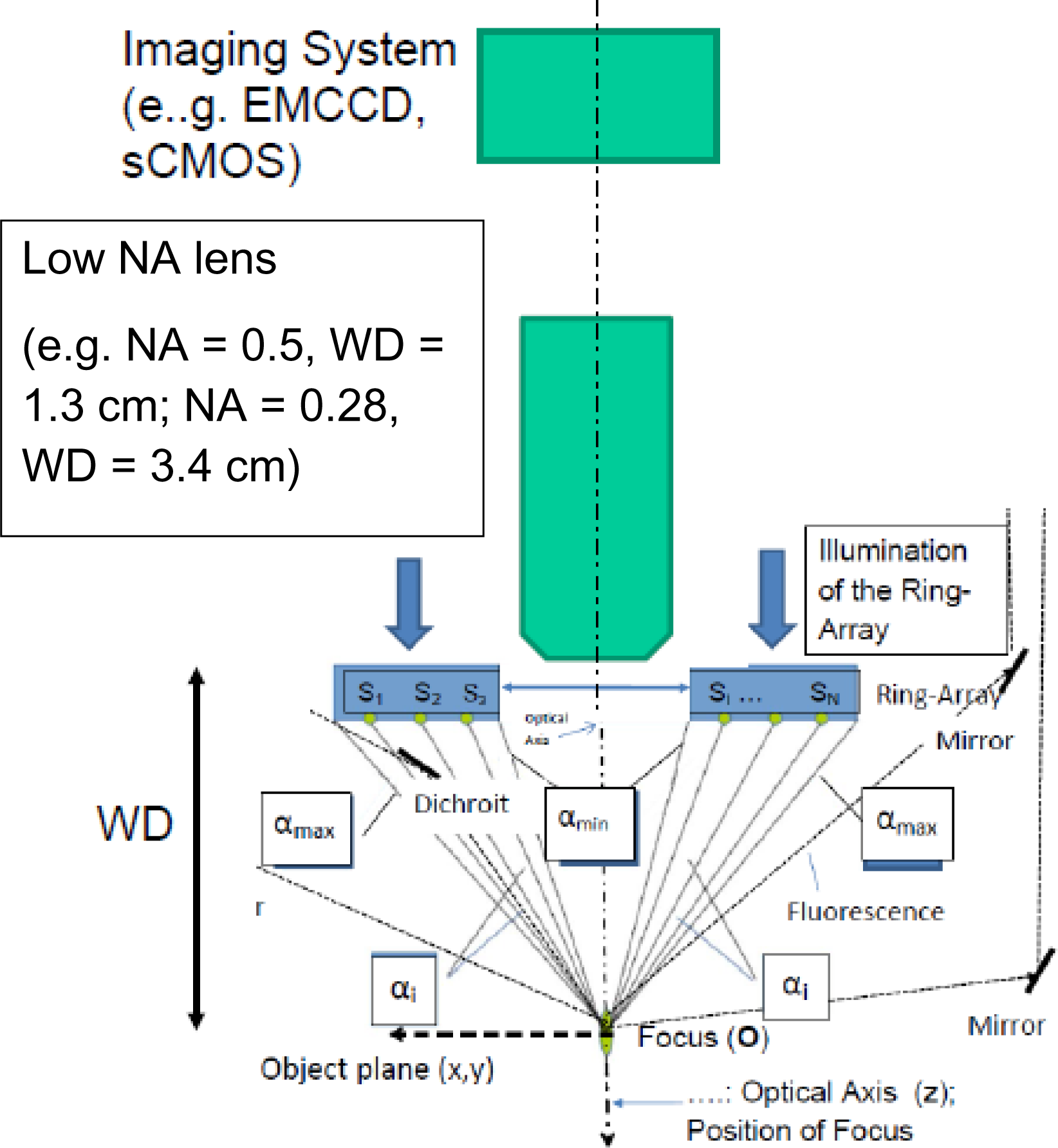
Combination of low NA/large working distance (WD) objective lens with Ring-Array illumination: The Interior Zone around the Optical Axis is the site of the low NA/Large WD objective lens. Structured or focal/donut illumination of selected sites is obtained by synthetic aperture illumination obtained by constructive interference from coherent sources arranged in a Ring-Array around the low NA objective lens. The focal/donut/SIM position is provided by the stage coordinates (x,y,z). In case of sufficient fluorescence emission, the detection of fluorescent targets can be performed by the low NA/Large WD objective lens; in addition, the fluorescence yield may be increased substantially by additional collection of photons using a single or more detectors, together with an appropriate arrangement of the Ring-Array sources. Altogether, a photon yield sufficient for nanometer range SRM resolution may be realized, using e.g. ultrastable, switchable nanographenes for labeling.

### General concept of “Ring-Array microscopy”

In the following, a general concept is described which makes it possible, by means of a suitable “ring-shaped” spatial distribution of coherent light sources (“Ring-Array”) with specifically adapted relationships of phase, polarization, direction and intensity between the coherent laser beams emitted from the light sources of the Ring-Array to realizeat extremely large working distances in the object plane illumination patterns (focused, donut-like, or structured) which allow a scanning based resolution corresponding to a high synthetic Numerical Aperture NA (for a summary of the principle rationale see Figure S1).In contrast to earlier concepts of synthetic aperture based enhanced resolution microscopy (Birk et al., 2017), the Ring-Array concept presented here is designed in such a way that an interior zone with a diameter D_interior_ (Figures. 1 – 3, 8) is left free of illumination sources S_i_. This makes possible to use this zone for other equipment required for direct correlative microscopy purposes. In addition, in contrast to earlier synthetic aperture configuration (Birk et al., 2017) requiring a three-dimensional arrangement of coherent illumination sources, the present Ring-Array design may be flat, i.e. all sources S_i_ may be located in a plane. Fig. 4 shows a schematic example for the distribution of coherent light sources S_i_ in such a Ring-Array configuration. For the numerical calculation of intensity, phase, direction of polarization and direction of propagation of the waves emitted by the sources S_i_ to implement the arrangement of the sources according to the Ring-Array concept and to determine their properties, it is necessary to specify the position of the sources in the Ring-Array in more detail. This can be done e.g. by polar coordinates, or by any other mapping procedures. The focal illumination distribution (volume indicated by the ellipse around **O** in Figs. (1-3) is determined by the constructive interference of the total of N coherent - e.g. collimated - rays S_i_ generated (for details of calculation see [von Hase et al., 2022]). When using the Ring-Array illumination according to the concept, maximum angles α_max_ =70° can be achieved; this corresponds to a synthetic numerical aperture (NA) of a lens of 0.94 (vacuum, n= 1); in contrast to such lens-based objectives with only a small working distance (typically in the range of 0.2 mm), the working distance (WD) in the illumination ring-array arrangement can in principle be chosen to have “any value”, e.g. WD = 5 cm. (see examples Figs. 5 – 7).

**Table 1:** Focused Ring-Array-Mode.

| **FWHM<sub>x</sub>** | **FWHM<sub>y</sub>** | **FWHM<sub>z</sub>** |
| --- | --- | --- |
| 0.497 $\lambda_{\text{exc}}$ | 0.599 $\lambda_{\text{exc}}$ | 1.355 $\lambda_{\text{exc}}$ |

1b) Focus size in the interior of a transparent material (ice, $n = 1.31$ ):
| **FWHM<sub>x</sub>** | **FWHM<sub>y</sub>** | **FWHM<sub>z</sub>** |
| --- | --- | --- |
| 0.489 $\lambda_{\text{exc}}$ | 0.727 $\lambda_{\text{exc}}$ | 1.361 $\lambda_{\text{exc}}$ |

| **FWHM<sub>x</sub>** | **FWHM<sub>y</sub>** | **FWHM<sub>z</sub>** |
| --- | --- | --- |
| 0.393 $\lambda_{\text{exc}}$ | 0.771 $\lambda_{\text{exc}}$ | 2.547 $\lambda_{\text{exc}}$ |

2b) Focal size within a transparent material (ice, $n = 1.31$ ):
| **FWHM<sub>x</sub>** | **FWHM<sub>y</sub>** | **FWHM<sub>z</sub>** |
| --- | --- | --- |
| 0.407 $\lambda_{\text{exc}}$ | 0.539 $\lambda_{\text{exc}}$ | 2.545 $\lambda_{\text{exc}}$ |

When conditions are applied according to confocal laser scanning fluorescence microscopy (CLSM), the achievable resolution is given by $\text{FWHM}_{xy}/\sqrt{2}$ .

**Fig. 4:**
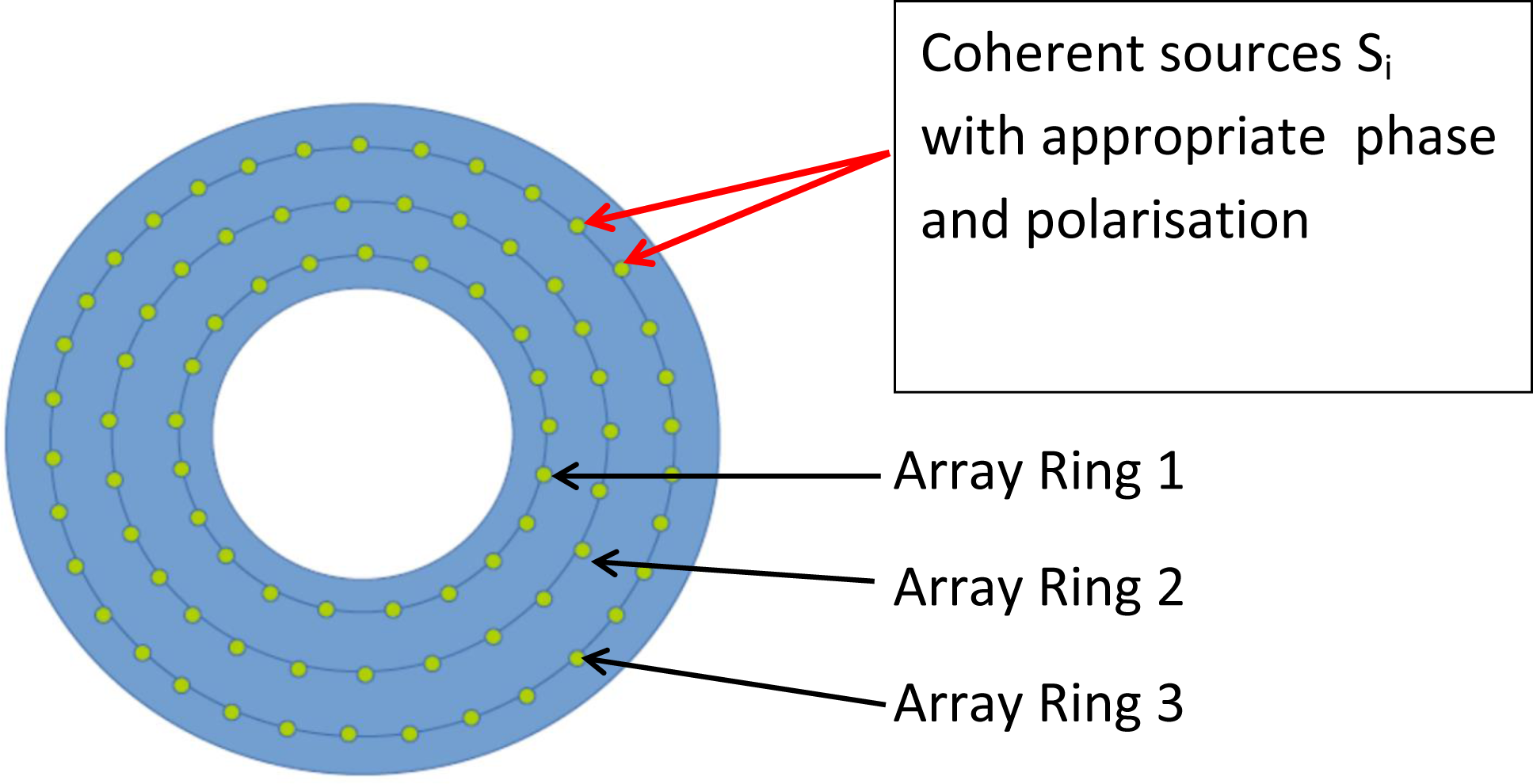
Schematic arrangement of the coherent sources Si within the Ring-Array. In this schematic implementation example, is is assumed that the S_i_ sources are arranged along three circular lines (“array rings”) at equidistant positions from each other; instead of 3 array rings, it can also be 1, 2, 4, 8 or more. Instead of arranging the emission sources S_i_ (S_1_,… S_N_) in concentric array rings, these can also be positioned in any other geometry given by the steric angle Ω_array_ (Fig. 2), provided that the values for phase, polarisation, intensity and direction of propagation of coherent waves emitted from the sources S_i_ are configured in such a way that the desired intensity distributions (see examples Figs. 5 – 7; Tables 1,2) can be obtained.

The numerical simulations indicated that in this way, in a FIB-SEM, or in other correlative microscope systems with the requirement of a very large working distance, at selected sites an optical resolution may be achieved which in the confocal scanning microscopy mode almost corresponds to the best optical resolution values with objective lenses of high NA, i.e. about 250 nm laterally and 900 nm axially (e.g., Figs. 6, 7; Table 1). With appropriately designed “Airy disc” confocal detection systems, this resolution might be further enhanced (Gardill et al., 2022).

### Nanosizing mode

The light microscopic measurement of the size of optically isolated nanostructures in the nm range has found a variety of biomedical applications (Cremer & Birk, 2022). Presently, however, so far only objective lenses with high NA and correspondingly low working distance and field of view have been used. Hence the application of such methods to correlative microscopy with large fields of view (to scan e.g. extended tissue sections, or nanostructures located in thick transparent specimen; or to analyse nanostructures in the vacuum chamber of a Focused Ion Beam Electron Microscope) has been regarded as impractical. This problem might be solved, however, by a special implementation of the Ring-Array Microscopy concept (Ring-Array-Nanosizing). Such systems may be applied wherever correlative microscopy based nanosizing of small objects in large fields of view at large working distances is required, e.g. for the fast image analysis of the transcriptional status of specific genes in large cell aggregates (Suppl. Material, Fig. S4); or in organoids a chip (Park et al., 2019), spheroids, or other, sufficiently transparent thick structures.

Using pairs of counter-propagating waves of opposite sources S_i_ in the Ring-Array (Supplementary Figure S2), structured illumination patterns may be realized at extremely large working distances, such as obtained in Standing Wave Field Microscopy (Bailey et al., 1993), or in Spatially Modulated Illumination (SMI) microscopy (Hausmann et al., 1997; Schneider et al., 1998; Albrecht et al., 2002; Failla et al. 2002a,b, 2003; Baddeley et al., 2007). Under the conditions of Ring-Array illumination (e.g. λ_exc_ = 488 nm, n = 1, θ_max_ = 70°, where θ_max_ the maximum angle between the directions of the two interfering ring-array beams to the optical axis), smallest distances between neigboring interference maxima (fringes) along the optical axis of Δ_min_(z) = λ_exc_/(2n cosθ_max_) = 713 nm would be obtained. Accordingly, along the object plane Δ_min_(xy) = λ_exc_/(2nsinθ_max_) = 260 nm is obtained for the fringe distance, and FWHM_fringe_ = λ_exc_/(4nsinθ_max_) = 130 nm for the fringe halfwidth. Extensive virtual microscopy calculations based on [Failla et al. 2002b] indicate that these conditions should still allow to determine by Ring-Array based “Nanosizing” the diameter and shape of a small optically isolated object with constant fluorescence emission down to diameters of < 100 nm in the direction of the beam propagation along the object plane, even at working distances of several cm (for further details see Supplementary Information, Figs. S2 – S4).

### Resolution enhancement down to the nanometer range

With additional use of donut intensity distributions in the Ring-Array “STED” mode (e.g., Figs. 6; Table 2; Fig. S6), even at a working distance up to several cm, an optical (object plane) resolution should be achievable down to the few tens of nm range; down to the 5 nm range in the Ring-Array “SMLM” mode (in combination with coherent fluorescence collection modes [Hell et al., 1994b]) approximately 1/100 λ_exc_; and down to the 1 nm range (approximately 1/500 λ_exc_) in the “MINFLUX/SIMFLUX” mode (Best et al., 2014; Balzerotti et al., 2017; Cnossen et al., 2020; Cremer & Birk, 2022).

**Tabele 2:** Donut Modus (see Figs. 6, S6)

| **D** | **d** | **$\Delta z_{\text{max}}$** |
| --- | --- | --- |
| 0.385 $\lambda_{\text{exc}}$ | 0.206 $\lambda_{\text{exc}}$ | 1.449 $\lambda_{\text{exc}}$ |

b) Donut intensity distributions in a transparent material (ice, $n = 1,31$ ):
| **D** | **d** | **$\Delta z_{\text{max}}$** |
| --- | --- | --- |
| 0.385 $\lambda_{\text{exc}}$ | 0.206 $\lambda_{\text{exc}}$ | 1.451 $\lambda_{\text{exc}}$ |

**2) $45^\circ < \text{aperture angle} < 70^\circ$ (For comparison with a lens based scanning microscope)** a) Donut intensity distributions in vacuum/air ( $n = 1$ ):
| **D** | **d** | **$\Delta z_{\text{max}}$** |
| --- | --- | --- |
| 0.343 $\lambda_{\text{exc}}$ | 0.356 $\lambda_{\text{exc}}$ | 2.549 $\lambda_{\text{exc}}$ |

b) Donut intensity distributions in a transparent material (ice, $n = 1,31$ ):
| **D** | **d** | **$\Delta z_{\text{max}}$** |
| --- | --- | --- |
| 0.343 $\lambda_{\text{exc}}$ | 0.354 $\lambda_{\text{exc}}$ | 2.551 $\lambda_{\text{exc}}$ |

To realize such enhanced resolution estimates, the collection of a sufficient number of photons is necessary. Fig. 8 shows a very schematic design how to do this. The total signal S_Fluor (x,y,z_), which is essential for imaging, results from the sum of the fluorescence signals F(x,y,z) measured by the individual detectors; by suitable optical elements/mirror constructions it is also possible to perform the fluorescence detection with only one detector. The image obtained by scanning the object at selected sites is formed from the totality of the signals measured for signals obtained for each individual object point. When using confocal detection arrays, the axial discrimination can be performed by means of of appropriately dimensioned pinholes.

For example, let us consider a Ring-Array illumination where object points are successively scanned and excited to fluorescence emission, assuming where, α_min_ = 45°, n = 1 (i.e. vacuum/air), and Ω_stage_ = 2π, the possible efficiency of the fluorescence emission collection corresponds to that achievable with an objective of numerical aperture 0.95 (Supplementary Fig. S5). In this case, the fluorescence emission obtained for a given fluorescence emission may be compared with an immersion objective with a numerical aperture of NA = 1.4: Under these assumptions, the fluorescence yield of the Ring-Array mode would be reduced by a factor of (1.4/0.95)^2^ = 2.2 x only; accordingly, the optical SMLM resolution would be reduced reduced by a factor of sqrt (2.2) = 1.4/0.95 = 1.5 x, assuming the same amount of registered photons.

Using in a Ring-Array configuration (Fig. 3) for photon collection the central low NA objective lens only, for NA= 0.1(WD = 5 cm), a photon yield about (1.4/0.1)^2^ ∼ 200x smaller than in present SMLM applications with high NA objective lenses is expected. For NA = 0.28 (WD = 3.4 cm), the fluorescence yield collected by such a lens is reduced by a factor (1.4/0.28)^2^ = 25 compared to a high NA immersion lens. Assuming e.g. at NA = 1.4 a collection rate of 5,000 photons of each “blinking” event (Liu et al., 2020) is achieved, 200 photons registered by a 0.28 lens alone may yield a theoretical localization precision (Cremer et al., 2017) of [0.51 x 488 nm]/[NA x (200)^0.5^] = 63 nm, or an opticl resolution around 150 nm. This problem may be solved, however, by additional photon collection measures (Supplementary Material, Fig. S5). In this way, up to ca. 50% of the photon yield typical for high NA SMLM may be obtained. Hence, in case of a coherent fluorescence collection scheme (Hell et al., 1994b) at the same amount of fluorescence excitation, the principle limit of resolution should be dimished only by a factor of sqrt(2) compared to a high NA immersion lens. Using novel fluorescence labels like nanographenes (Liu et al. 2020), the adverse effects of fluorescence collection may even be compensated. For example, several types of nanographenes (Liu et al., 2020) have a high brightness of more than 5,000 (detected photons per fluorescent on event) using a 1.4 high NA objective lens, in addition providing higher photostabilities as compared to other fluophores.

From the technical point of view, coherent detection is challenging to realize. However, even if due to lack of coherent detection the collection of photons is restricted to increase the total amount of registered photons only, this may still be highly useful, e.g. for spectroscopic analysis of single molecules in a very large field of view and correspondingly large working distances.

With Ring-Array illumination microscopy allowing very large working distances and enhanced resolution, the entire process of preparing objects of interest described above may be performed directly in the FIB-SEM (dCLEM), or in a low NA optical screening system (dCOLM). This has the advantage that 1) sample transfer is avoided; 2) in dCLEM, the ion beam can be guided under light-optical control, greatly reducing sample-damaging electron illumination; in dCOLM, no mechanical modifications of the system are required for switching the microscopy mode; and 3) with Ring-Array illumination microscopy using coherent light according to the concept (Figs. 1 - 3; 8), the site can be located more precisely, even 3-dimensionally; using structured illumination modes, the size of optically isolated objects may be estimated in 3D down to sizes ≤ 100 nm. Novel fluorochromes, based e.g. on buffer independent nanographenes (Liu et al., 2020), can also be used to localize sites. This saves a considerable amount of time and allows one to analyze exactly the target site that is important.

The concept of Ring-Array microscopy described here will not be limited to cryo-FIB-SEM applications, but may revolutionize the entire volume microscopy: For example, samples labeled with nanographenes or colloidal gold are prepared for electron microscopy, and bands of serial thin sections are mounted on an electrically conductive support. These are placed in a field emission grid EM equipped with the ring array illumination of the invention and associated facilities for fluorescence excitation and detection. Then the light and electron microscopic images are taken of each section. This direct application is characterized not only by significant time savings, but also by the fact that no changes, shrinkage, distortion, etc. of the sections occur between the light microscopic and electron microscopic images. Since, for example, nanographenes and colloidal gold particles retain their fluorescence and “blink” behavior even after considerable chemical treatment, the “ring array microscopy” according to the invention can be applied to a large variety of sample preparation techniques.

In the Ring-Array illumination microscopy method presented here, due to the possibility of individually controlling the characteristics (e.g., intensity, phase, polarization state, propagation direction, divergence) of a finite number of coherent beams (e.g. N = 190, or N =760) individually, in addition to point-scanning features corresponding to a confocal microscope type (Cremer & Cremer, 1978), a number of other implementations are possible, such as STED, SIM, SMLM, MINFLUX, or SIMFLUX microscopy, light sheet microscopy (“lightsheet”), optical projection microscopy, or axial tomography microscopy (Staier et al., 2011; Schneckenburger et al., 2020).

Instead of being used for construction elements of a FIB-SEM, the inner zone free of excitation light beams S_i_ (solid angle Ω_center_, Fig. 2) with the diameter D_interior_ can also be used in other microscopy applications where high light-optical resolution together with a large working distance WD is to be achieved, such as X-ray microscopy, or other methods of ultrahigh-resolution microscopy using particle radiation or high-energy photon radiation.

Further advantageous applications of the Ring-Array concept also arise in certain applications of light microscopy where large working distances are required or desirable. This is the case, for example, in stereomicroscopy, or in material analysis investigations, e.g. the light-optical inspection of electronic components. For example, in this case the inner zone of the ring array shown in Figs. 1-3, 8 (associated spatial angle Ω_center_) as well as the area delineated by the spatial angle Ω_interior_ can be used for a lens-based objective lens of low NA but large working distance and correspondingly large field of view; this allows to localize object regions of interest (ROIs) to be analyzed in more detail, first at correspondingly low optical resolution, which are then examined in more detail with the aid of the Ring-Array illumination method, in conjunction with a scanning microscopy method at greatly enhanced resolution (e.g. using the STED, SIM, SMLM, MINFLUX/SIMFLUX mode). Such a combination would be suitable, for example, to significantly speed up microscopic procedures in which relatively few localized objects (ROIs) need to be analyzed at high resolution in large fields of view. Examples would be the analysis of nanostructures of selected cells in tissues (Oleksiuk et al. 2015; Neumann et al. 2020; Cremer et al. 2017, 2020; Lang et al. 2021); the analysis of pathogenic viruses or bacteria (Cremer 2011; Cremer et al., 2014); or the analysis of nanostructural changes in surfaces (Liu et al. 2019). In addition, the Ring-Array Microsopy approach may also be used to create 3D data sets of large volumes using the axial tomography technique (Staier et al., 2011; Richter et al., 2017; Schneckenburger et al., 2020). In this method, a sufficiently transparent and optically homogeneous object is rotated by specific angles; at each angle, a microscope image is registered; the series of images is then used to computationally reconstruct a 3D image at an enhanced isotropic resolution (Heintzmann & Cremer, 2002), corresponding to the object plane resolution achieved by the special Ring-Array mode. For example, for a NA = 0.28/WD = 3.4 cm lens (n = 1, λ_exc_ = 488 nm) and an assumed Ring Array object plane resolution of 300 nm (focused mode, Fig. 6; Table 1), this would result in an enhancement of the 3D resolution (measured by V_obs_, using eq. (1), (2a) for lateral/axial resolution) of a factor ∼ 23x, compared to a conventional microscope setup with a NA = 0.28 lens.

The examples of ring-array illumination arrangements given are based on the assumption that the coherent light sources used to generate an appropriately small scanning focus (or other intensity distributions suitable for methods of scanning light microscopy, such as toroidal (“donut” shaped intensity patterns) in the object plane are typically uniformly distributed with respect to the angle between two adjacent sources within the ring-array (Fig. 4). (i.e., within a solid angle Ω_array_ (Figs. 1,2, 8) with suitable maximum aperture, e.g., Ω_arraymax_ = 1.3 π, corresponding to a NA in vacuum (n = 1) of 0.94. Instead of a uniform arrangement of the sources as in the example calculations given here (Figs. 5-7), other distributions of the sources in the Ring-Array arrangement are also possible. Coherence of the radiation emitted by the light sources is achieved by suitable illumination of the ring array with coherent light (in particular, e.g., Laser radiation of suitable wavelength and intensity).

**Fig. 5:**
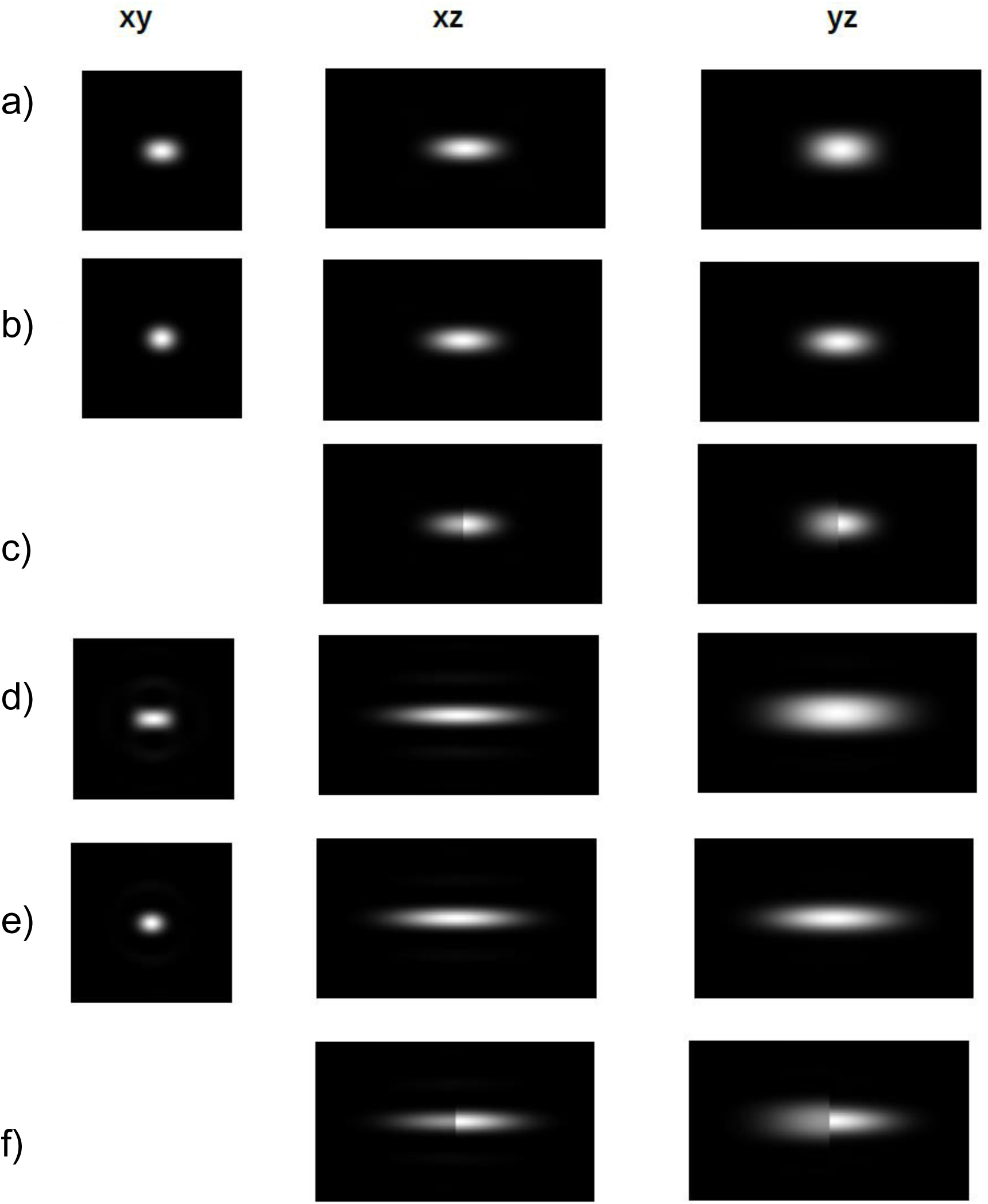
Example images for the numerical calculation of fused intensity distributions in the object plane using a Ring-Array configuration. For the numerical calculation of object plane illumination patterns generated by Ring-Array, configurations with α_min_> 0, N = 190 sources (S_1_,, S_2_,..S_190_) of the same wavelength λ_exc_ and radiant power were assumed. These sources were uniformly arranged in 4 array rings (see Fig.4), where the average angular separation Δα between two adjacent source positions was 6°, and the angle from the aperture edges to the adjacent rings (Fig. 1,2) was Δα/2 = 3°. An equal radiant power was assumed for all individual waves. For comparison with the result that would be expected for a lens with a numerical aperture NA = nsin(α_max_), the same number of sources was uniformly distributed in a source array with α_max_ = 70° but α_min_ = 0 (Figs. 5a-c). xy: intensity distribution in the object plane (plane orthogonal to the optical axis, see Fig.1. xz, yz: intensity distributions (xz, yz sections) along the optical axis (z), see Fig. 1. The numerical calculation was performed for a large working distance WD = L_Array_, e.g. WD = 5 cm. For further details see Table 1. _a)_ Refractive index n = 1 (air, vacuum), with a half-aperture angle (Fig. 1) α_max_ = 70° and α_min_ = 0°, i.e. Ω_center_ = 0 sr. The numerical simulations for this setup correspond to an objective lens with a numerical aperture NA = 1 x sin (70°) = 0.94; b) Conditions as in a) with the difference that the focal area is completely inside an ice cuvette with a refractive index n = 1.31; c) Conditions as in a) with the difference that the surface of the ice cuvette is placed at the height of the focus center (**O**); d) Refractive index n = 1 (air, vacuum), with a half-opening angle (Fig. 1) α_max_ = 70° and α_min_ = 45.5°. The numerical simulations for this arrangement correspond to a ring-array arrangement with a D_interior_ (Figs. 1, 2) corresponding to a solid angle Ω_center_ = 0.6π sr; (e) Focal intensity distribution with complete “immersion” of the focal area into a transparennt material (e.g. ice) with a refraction index n = 1.31 and a half aperture angle (Fig. 1) α_max_ = 70° and α_min_ = 45.5°. The numerical simulations for this arrangement correspond to a specific Ring-Array arrangement with a D_interior_ (Figs. 1, 2) corresponding to a solid angle Ω_center_ = 0.6π sr. (f) Focal intensity distribution at the surface of the ice (transition from n = 1 to n = 1.31. Ring-Array illumination with α_max_ = 70° and α_min_ = 45.5°. The numerical simulation shows the intensity distribution in the direction of the optical axis in the ice (right part of the image) compared to the vacuum region (left part of the image).

In contrast to “ring light microscopy” techniques as used e.g. in stereo microscopy for wide-field illumination using LEDs, the light sources S_i_ located in the Ring-Array emit waves with very specific mutual phase relationships, propagation directions, polarization states and powers due to the coherent illumination of the ring array, which can also be individually adjusted in such a way that very specific, diffraction-limited local intensity distributions are generated in the object plane, which, using scanning microscopy techniques, enable a high-resolution and/or super-resolution light microscopy at large working distances directly in a focused ion beam scanning electron microscope (FIB-SEM), or another ultra-high resolution microscope system based on particle radiation or X-rays, or another microscope system with the requirement of large working distances.

Optical fiber techniques may be one of the technical solutions to experimentally realize the coherent Ring-Array illumination needed, since it provides more flexible, feasible and robust laser beam systems. Currently, with suitable options of optical fiber components (such as polarization maintaining fibers, fiber polarizer, fiber attenaturator, fiber phase control, integrated beam collimators for fiber optics etc.), the beam properties (e.g. propagation directions, phase, power and polarization) can all be tuned.

In order to realize the “point-by-point” scanning of the object required when using the Ring-Array illumination method described herein for focused/donut imaging, “stage scanning” (movement of the object) according to the state of the art (e.g., Cremer & Cremer 1978; Hell et al. 1994; Hänninen et al., 1995) is assumed in the example applications discussed. However, according to the general features of the concept, “beam scanning” (e.g., Hell et al., 1999) by suitable periodic movements of the Ring-Array might also be possible. Hereby, the Ring-Array microscopy allows, in addition to a very large working distance, a substantially enhanced optical resolution, which is NOT possible with the previous “ring-light” microscopy; due to the lack of mutual coherence of the waves emitted by the “ring-light” array, this latter technique only allows an improved wide-field illumination, while the optical resolution is given by the numerical aperture of the objective used according to Eq.(1,2); for example, for λ_exc_ = 488 nm and a numerical aperture NA = 0.15 (n = 1) typical for stereomicroscopy, a working distance of 6 cm results in a lateral optical resolution of about 1.7 µm and an axial one of about 22 µm (according to relation 2a), and 20 µm (eq. 2b), respectively. In contrast, using the coherent ring array microscopy approach at the same working distance (6 cm) in focused mode (e.g., Figs. 5 -7; Table 1) in conjunction with confocal laser scanning fluorescence microscopy (CLSM), a lateral resolution that is approximately 5x enhanced and an axial resolution that is approximately 18x enhanced than achievable with an objective lens with NA = 0.15 (n = 1, λ_exc_ = 488 nm) is predicted. The volume resolution (observation volume according to Eq. (3); Table 1) can even be improved by a factor of ∼ 135x; e.g., the observation volume important for finding the object region of interest during investigations in a FIB-SEM (or in another microscopy system with a large working distance) can thus be reduced by this amount and hence substantially simplified. When using the Ring-Array illumination method in conjunction with super-resolution implementations such as STED, SMLM, MINFLUX, or SIMFLUX (e.g., using toroidal/donut intensity distributions, see Figs. 6, S6; Table 2), the resolution of the Ring-Array microscopy method can be further enhanced to the molecular resolution range while maintaining large working distances (e.g., 4 cm, 5 cm, or even 10 cm).

**Fig. 6:**
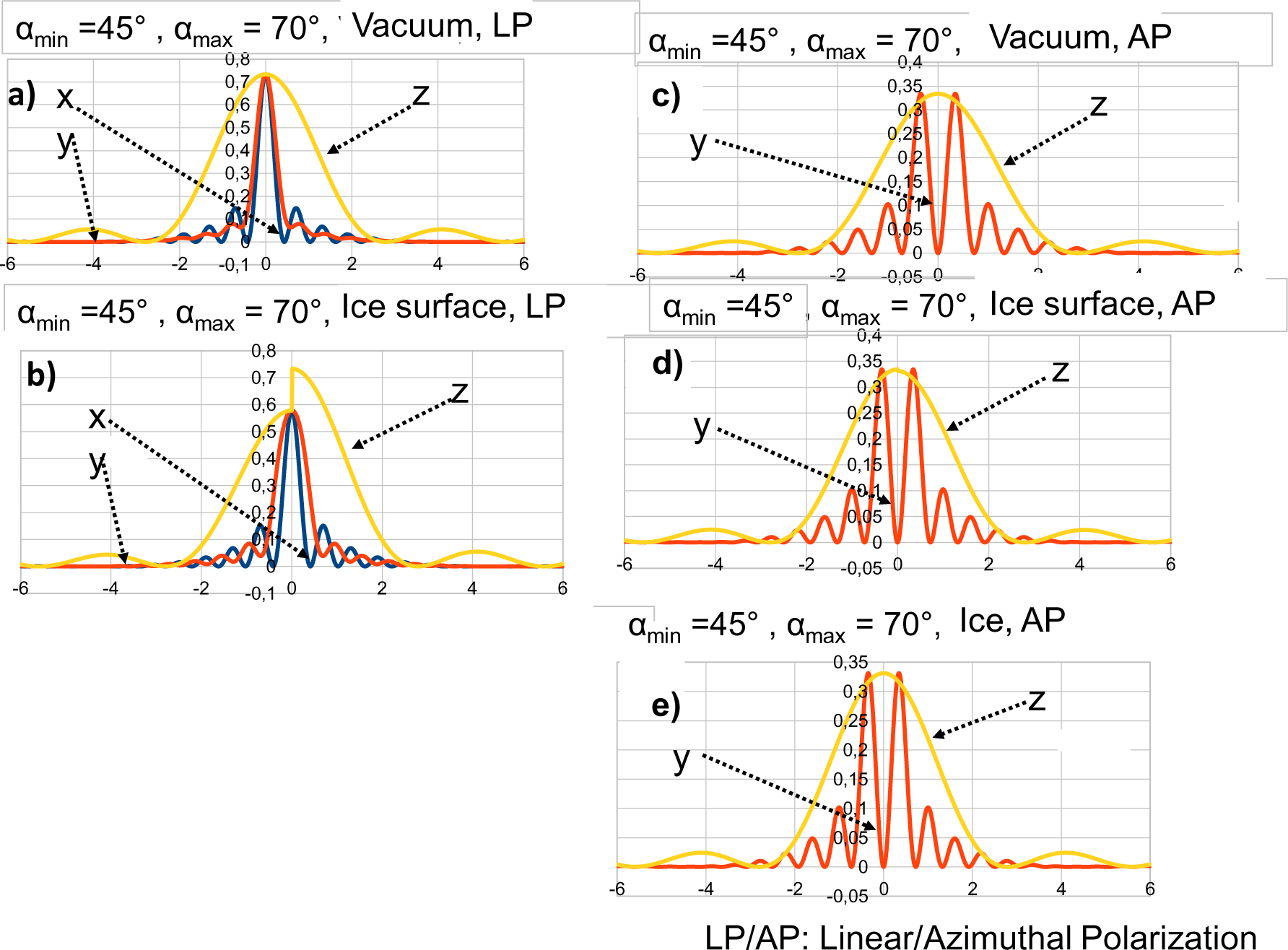
Illumination Ring-Array generated intensity distributions in the object plane. Based on the positioning of the coherent sources S_i_ (Figs. 1-3) in the illumination Ring-Array arrangement (Fig. 4), the resulting focal or toroidal/donut intensity distributions in the object plane through the focal center **O** (Fig. 1,2,8) were calculated, using procedures according to the state of the art (Feynman, R.2006; Birk et al., 2017; von Hase et al., 2022). Simulation conditions: Number of sources S_N_: N = 190 (linear polarization for cases a) - b); azimuthal polarization for cases c) - e); in each case a uniform distribution of the S_N_ = 190 sources in 4 rings (Fig. 4) was assumed; angular difference Δα between the rays (wave vectors) of two adjacent sources: 6°. α_min_ = 45°; α_max_ = 70°. Using a radial symmetry, this results in the following values for the corresponding solid angles (n = 1): Ω_center_ = 2π x [1 - cos(45°)] = 1.84 sr; and Ω_array_ = 2π x [1 - cos(70°)] - 1.84 sr = 4.13 sr - 1.84 sr = 2.29 sr. In ice (n = 1.31), values for the half-opening angle range between β_min_ = 33° and β_max_ = 45.8° are obtained. a), b): Linear polarization (LP) of the constructively interfering beams emitted from the Ring-Array sources S_i_ results in focused intensity distributions around the center **O** (Fig.1,2). a) refraction index n = 1 (vacuum, air) in the entire focal region was assumed; b) Intensity distribution around the focal region **O** at the boundary beween n = 1 and transparent material (n = 1.31, e.g. ice); c-e) Azimuthal polarization (AP) of the constructively interfering beams emitted from the Ring-Array sources S_i_ results in donut shaped intensity distributions around the center **O**. c) refraction index n = 1 (vacuum, air) in the entire focal region was assumed; d) Intensity distribution around the focal region **O** at the boundary beween n = 1 and transparent material (n = 1.31, e.g.ice); e) Intensity distribution around the focal region O positioned inside a transparent material (n = 1.31, e.g. ice). Ordinate: Intensity (arbitrary units). Abscissa: Distance (unit in vacuum wavelengths λ_exc_) from the center of focus **O** (xyz = 0, Fig. 1) in xy direction (object plane), or in the direction (z) of the optical axis, respectively; the same unit applies to vacuum wavelengths for STED/MINFLUX excitation. Intensity in x-direction is marked in blue, in y-direction marked in red; the intensity in z-direction (along optical axis) is marked in yellow. The intensity distributions are given around the maximum of the focal intensity distribution and around the maxima of the donut distribution, respectively. If only the x- or only the y-direction is indicated, the intensity distributions for x- and y-coordinates are practically identical. Negative z-coordinates indicate a position below **O** (i.e. distance to illumination Ring-Array > L_Array_).

Due to the large working distances and available solid angles possible with the ring-array illumination method according to the invention, efficient detection of the signal induced in the object region with the Ring-Array illumination method (e.g., fluorescence excitation) is also greatly facilitated (Fig. 8; Fig.S5). For geometric reasons, Ω_array_ = 4π is not possible in a FIB SEM due to the instrumentation required for the object mounting and detection devices (see Fig. 2 in Gorelick et al. 2019). Similar restrictions are also obtained in other microscopy systems. Typically, the solid angle Ω_array_ in implementations of ring-array illumination according to the invention will be limited to values below π. Despite this limitation, however, local intensity distributions can be realized in the object plane by Ring-Array illumination according to the invention, allowing high resolution even at extremely large working distances.

By a suitable configuration of the coherent light sources of the Ring-Array illumination, it is furthermore possible to generate patterns of local intensity distributions (e.g. multiples of single maxima or torus-shaped distributions) in the object space, whose distances are larger than the half-widths (FWHM) of the respective intensity distributions (examples Figs. 7), and which are therefore suitable, with the aid of such patterns (e.g. Bingen et al. 2011; Chmyrov et al. 2013) to accelerate the speed of imaging quite substantially.

**Fig. 7:**
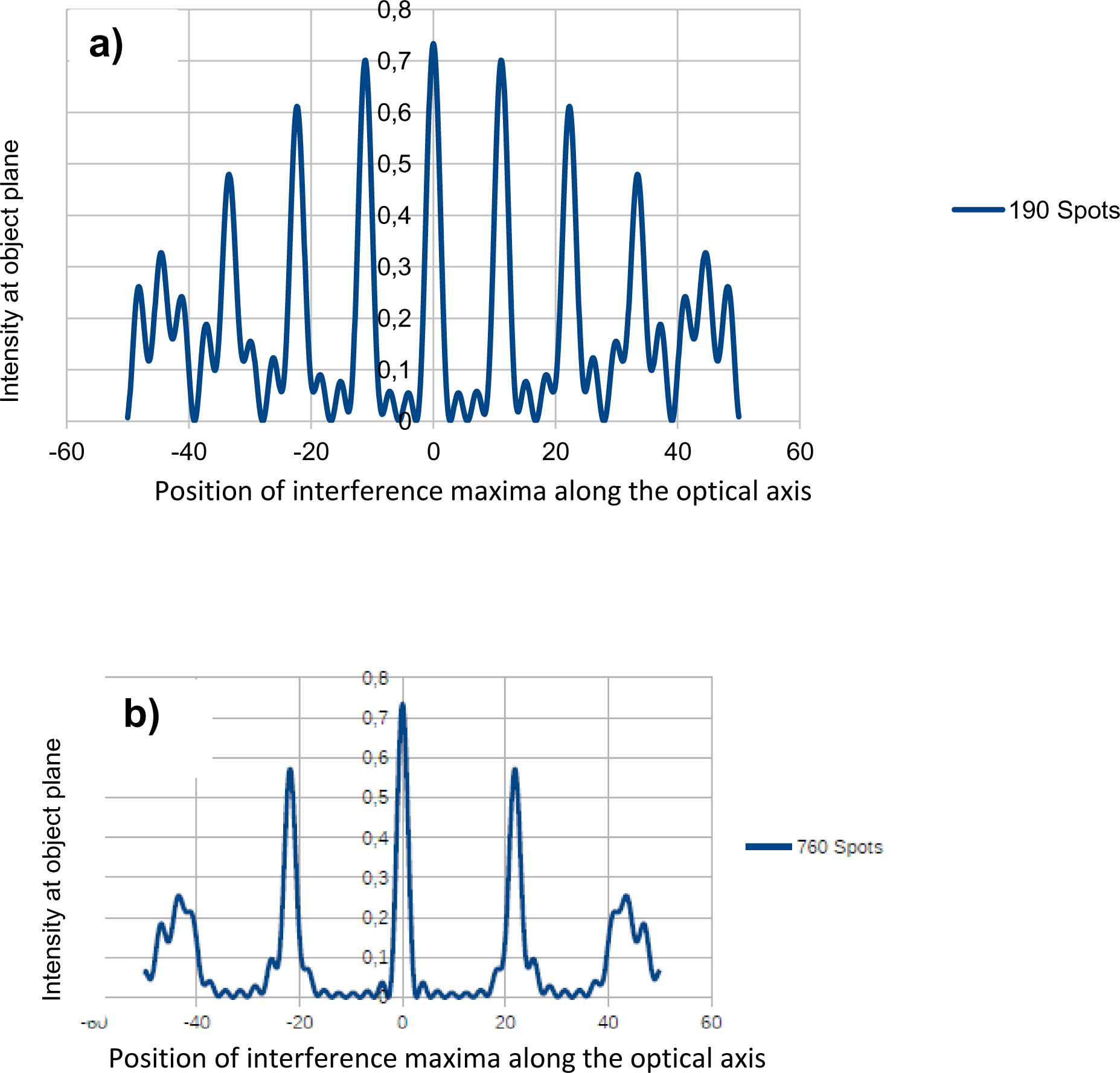
Ring-Array Intensity distribution along the optical axis (z) for S_N_ = 760 sources. a) Assumptions:α_min_ = 45°; α_max_ = 70°, N = 190 sources (“spots”) in 4 array rings; angular distance between the sources Δα = 6°; linear polarization of collimated waves. The intensity main maxima have a sufficiently large distance in the z-direction from each other (for λ_exc_ = 488 nm, e.g., about 5 µm to allow 3D imaging in confocal mode or by suitable deconvolution algorithms according to state-of-the-art (e.g., Hänninen et al. 1995). Periodic focal intensity main maxima at a distance greater than their half-width can be used to accelerate scanning microscopic analysis according to the analysis according to the state of the art. b) As in a), but for S_N_ = 760 sources (beams emitted from the Ring-Array); distance between the main maxima is increased to about 10 µm. α_max_ = 70°corresponds to a numerical aperture (NA) of an objective of 0.94 (vacuum/air, n= 1); however, in contrast to such lens-based objectives, the working distance (WD) in the illumination ring arrangement can in principle be chosen “arbitrarily large”, e.g. WD = 4 or 5 cm, or more.

The above stated theoretical and practical limitations of optical resolution at large working distances in an integrated FIB-SEM system (or other large working distance microscope systems) are due to the low numerical aperture of the objective lenses required for correlated light optical excitation and necessary to realize the large working distances required to fit, for example, the required light optics into the vacuum chamber of the FIB-SEM (for a typical arrangement, see e.g. [Gorelick et al. 2019]). In the Ring-Array Microscopy approach, these limitations are overcome, i.e., the optical resolution is substantially enhanced at large working distances, by using an illumination device with multiple coherent beams starting from an annular ring array arrangement and directed at a given object region with appropriate synthetic aperture, mutual phase relationships, polarizations, propagation directions, powers, and divergences (e.g., collimated), while not interfering with the other functions of, e.g., a FIB-SEM (dCLEM), or low NA screening (dCOLM). In the numerical example calculations described here (Figs. 6, Tables 1,2), collimated coherent beams were assumed; however, similar results are also possible with constructive interference of coherent spherical waves.

With the Ring-Array illumination principle, typically single molecule positions can be measured in SMLM mode in an object range of the desired size (e.g., diameter in the range 0.5 µm - 1 µm), with an optical resolution (smallest distance of the separately localized single molecules) down to the range of a few nanometers; with additional use of the MINFLUX option (Balzarotti et al., 2017; Gwosch et al., 2020), or with additional use of structured illumination (SIMFLUX, Best, 2014; Best et al., 2014; Cnossen et al., 2020), a resolution down to the 1 nm range (approximately 1/500 λ_exc_) can be achieved (Cremer & Birk, 2022). Since many target structures have diameters much smaller than 0.5 µm - 1 µm, the application of the Ring-Array illumination in the SMLM/MINFLUX/SIMFLUX mode should also be extremely advantageous: For example, a direct Ring-Array based combination of SMLM with MINFLUX/SIMFLUX/structured illumination makes it possible to analyze the molecular conformation of important proteins much more advantageously than is possible according to the state of the art (Weisenburger et al. 2017).

If single molecule microscopy/SMLM integrated in a FIB-SEM or into a low NA light optical system is to be performed with the aid of a larger illumination volume, this is also possible by suitably changing the phase, polarization, intensity and direction of the beams emitted by the ring-array arrangement according to the concept described. The same applies to the further improvement of the three-dimensional resolution with the help of e.g. torus-shaped intensity distributions along the optical axis for the realization of 3D-STED microscopy (Sahl & Hell, 2019).

The illumination pattern required for the combination of structured illumination and localization microscopy/SMLM (Best et al., 2014) can also be realized with the aid of the Ring-Array arrangement by a suitable combination of phase, intensity and utilized number of light sources of the Ring-Array.

### Experimental implementation

To experimentally realize Ring-Array microscopy, a large variety of approaches are feasible. A first option would be Ring-Array based Nanosizing (see above); since this requires only pairs of interfering illumination beams without phase matching, from the technical side its implementation should be straightforward. In the following, a few general problems are discussed to make possible also optical resolution enhancements down to the nm-range, using extremely large working distances. For more details, see Supplementary Material.

### Coherent multi-beam illumination

The interference of multiple beams used for the improvement of the optical resolution according to the principle of Ring-Array Microscopy, with simultaneous large working distance, may be realized by a suitably placed ring-shaped arrangement of e.g. collimated coherent beams; i.e. the beams emanating from the light sources S_i_ of the Ring-Arrays, which are generated by means of the illumination of the Ring-Array arrangement (Figs. 1 - 4, 8), have fixed phase relationships to each other, which can be individually determined by the details of the Ring-Array arrangement; the resulting coherent multi-beam illumination is performed using a Ring-Array zone that is free of FIB-SEM instrumentation or components required in other microscopy applications (“Ring-Array” arrangement, see example Figs. 4). This Ring-Array zone, containing the output points (light sources S_i_) for the coherent beams illuminating the object, covers a solid angle Ω_array_ (determined from the ring center, see Fig. 2; Fig. 8); The space containing the illumination and detection elements of e.g. a FIB-SEM covers a solid angle Ω_center_ and is free of coherent ring-array illumination beams (see Fig. 2). The space where the object holder of e.g. a FIB-SEM or other components are located covers a solid angle Ω_bottom_ and is also free of Ring-Array illumination beams. Therefore, the Ring-Array zone containing the multiple beams for ring array illumination covers a solid angle Ωarray = 4π - Ω_center_ - Ω_bottom_.

**Fig. 8:**
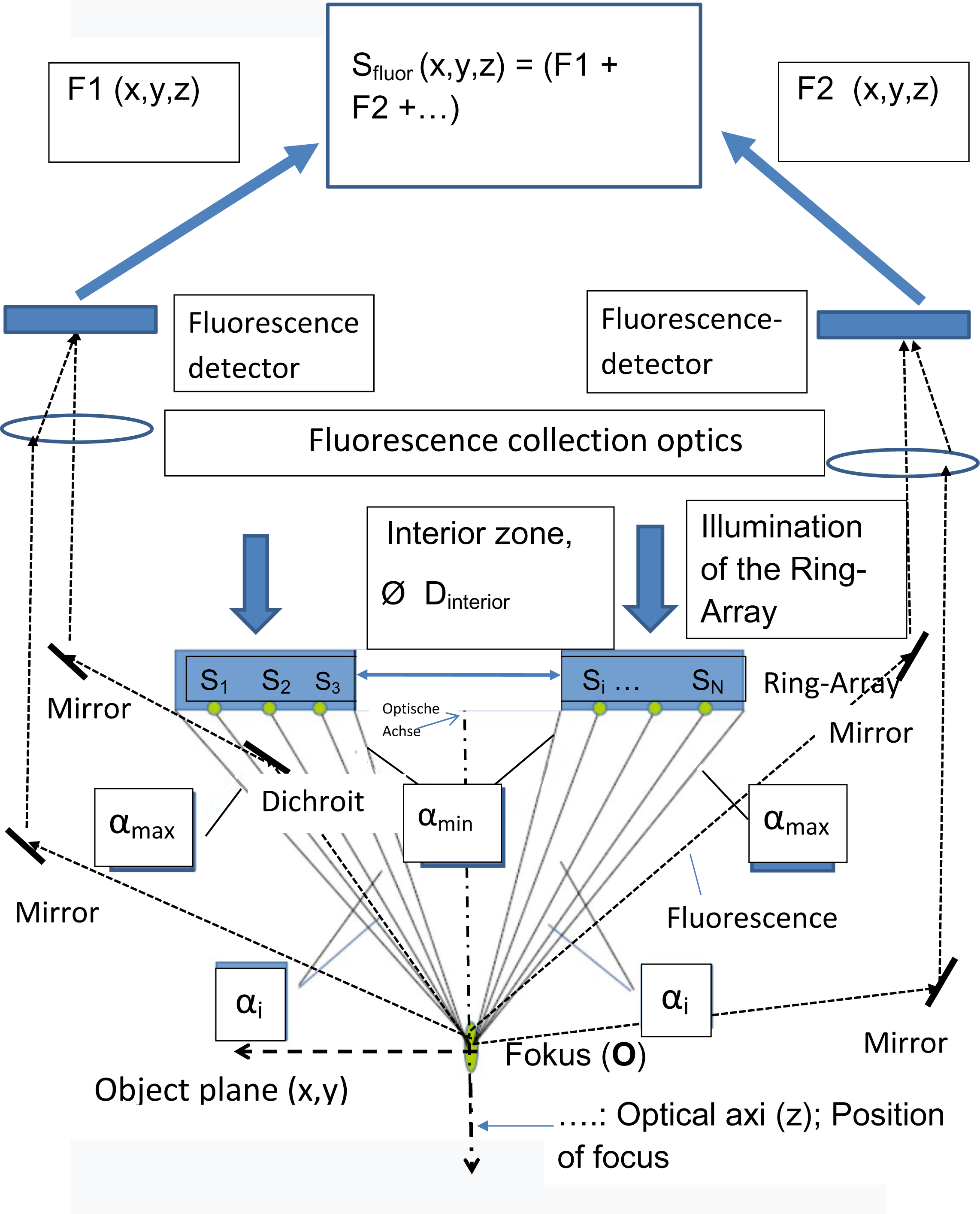
Example of the arrangement of a fluorescence detection system with high photon collection efficiency with localized excitation by the illumination ring array (schematic). The Ring-Array arrangement (see Figs. 1-3,4) makes it possible to collect the photons emitted from the object plane at a very large working distance with the help of suitable dichroitic mirrors (within Ω_array_) with maximum transmission for the excitation wavelength(s) and maximum reflection for the fluorescence emission of the object as well as with the help of one or more detectors (e.g. “point detectors” or area sensors (CCD, sCMOS). In the scanning mode, the positioning information is provided by the center of the focal/donut illumination pattern. The scheme is only intended to convey that an efficient photon collection is possible also at a very large WD. Many other arrangements are also feasible (see e.g. Suppl. Material, Fig. S5).

Ω_center_ and Ω_bottom_ depend on the exact configuration of the considered microscope system - e.g. a FIB-SEM, or a low NA screening system. Both Ω_center_ and Ω_bottom_ are expetced to have values substantially above zero, e.g. Ω_center_ > = 0.4 π and Ω_bottom_ > π. Thus, the Ring-Array arrangement is substantially different from previous illumination arrangements that provide for a homogeneous distribution of coherent light sources in an array surface corresponding to a complete spherical section around the optical axis (Birk et al., 2017), i.e., Ω_center_ = 0, in contrast to the arrangement described here, with Ω_center_ values typically much larger than zero (e.g., in the application examples described here, Ω_center_ = 2π[1 - cos (45°)] = 0.585π was assumed.

In addition to “point scanning” modes (including STED, MINFLUX/SIMFLUX, or single molecule microscopy techniques), the present concept of Ring-Array illumination can also be applied to enable pattern based scanning modes such as light sheet (“lightsheet”) microscopy, optical projection microscopy, or axial tomography at large working distances in an integrated FIB-SEM system (Schneckenburger et al., 2020).

In addition to its use in FIB-SEM systems, the Ring-Array illumination method can be used wherever obstacles inhibit illumination beam arrays or other components used for imaging that correspond to conventional lenses or lens systems, such as in stereo microscopes, or microscopes for non-contact material analysis at large working distances.

A major advantage of the Ring-Array illumination method described is the ring-shaped arrangement of the coherent light sources used to illuminate the object while maintaining a large working distance. For example, the inner diameter (d_interior_) of the ring-array arrangement (Figs. 1, 2, 3, 8) can be chosen in such a way that the components required for particle radiation and detection in a FIB-SEM (or in another microscope system) are enclosed by the Ring-Array.

In principle, all geometric shapes of the Ring-Array illuminatioin configuration are considered, provided they meet the general conditions of Ring-Array microscopy, e.g. oval, square, or rectangular requirements; the arrangement of the coherent light sources S_i_ (Figs. 1-3, 4, 8) also does not necessarily always have to be in a plane, but may also be within a volume.

In particular, flat designs of the Ring-Array allow a substantial simplification of the production of the underlying optical elements, e.g. by nanolithographic techniques (Sharma et al., 2022).

A key feature of the Ring-Array concept is the use of multiple coherent beams emanating from sources emitted over a specific solid angle Ω_array_ (see Fig. 2) within a typically flat ring zone in the “FIB-SEM space”, or more generally in the “illumination space” that is free of FIB-SEM instruments or other components, with the option to control the power, polarization, direction, and divergence of these coherent beams. As above-mentioned, such beam systems can be potentially realized by fiber optics in practical applications. Fiber optics based laser delivery systems are more flexible and have much less free space limitations. Meanwhile, the beam properties (power, polarization, direction and beam angles) can all be controlled by optical fiber based components which has similar/same properties as free space optics.

When integrated into other microscopy systems where Ring-Array illumination is advantageous, the “Inner Zone” in Figs. 1-3, 8 (indicated by the solid angle Ω_center_, Fig. 2) is occupied by design elements of such other microscopy systems instead of FIB-SEM components (e.g., suitable low NA but large working distance objective lenses such as in stereomicroscopy, or microscopic inspection of electronic components).

In the Ring-Array illumination concept, it is possible for a given source (S_i_) to be located anywhere along a line through the focus center **O** = (0,0,0) and the actual position (*xi*, *yi, zi*) of the source S_i_ (see Figs. 1, 2, 8); under this condition, the distances of the light sources (S_i_ to **O**) can theoretically be made “arbitrarily large” (e.g. 5 cm, 6 cm, 10 cm, or more), without changing the intensity distribution around the center at **O** = (0,0,0); Therefore, “arbitrarily large” working distances (distance from light sources S_i_ to an object located at or near **O**) can be realized without changing the intensity distribution at the focus center **O** (or more generally, without changing the point image function PSF of the illumination of a “point” fluorescent object located at O), in contrast to high NA objective lens based microscopy. The working distance WD (distance of the Ring-Array center to **O**, see Fig. 1) can be chosen in such a way that all elements needed in a FIB-SEM for particle/electron excitation as well as the associated detection optics are not affected in their function. The same applies to other microscopy systems in which Ring-Array illumination may be advantageous.For practical reasons, the working distance is likely not to exceed 5 -10 cm.

The values for the half-widths (FWHM) obtained on the basis of numerical simulations of focal intensity distributions for an excitation wavelength of λ_exc_ = 488 nm when Ring arrays are used according to the invention are summarized in Table 1 (see also Figs. 5, 6ab; 7), and in Table 2 for torus-shaped intensity distributions (see also Figs. 6c-e; Fig. S6). They show that both focused mode (Table 1) and STED/MINFLUX mode (Table 2) can produce focal diameter or torus (“donut”) configurations in the ice, even at very large working distances, equivalent to those obtained with lens-based microscope systems at high numerical aperture (but very small working distances not useful for FIB-SEM and many other microscopy applications).

As an example for the focal diameters/donut dimensions achievable according to Table 1, 2 and units of λ_exc_ given there (equally valid e.g. for λ_STED_, λ_MINFLUX_ or λ_SIMFLUX_), let us assume λ_exc_ = 488 nm; a typical focal size (FWHM_xy_) of about 0.6 λ_exc_ as well as 2.5λ_exc_ (z) then corresponds to approx. 290 nm in the object plane (xy) and 1.2 µm along the optical axis (z), respectively; this is sufficient for coarse screening of a sample for initial containment of objects of interest. The values given in Table 2 for the torus mode (STED/MINFLUX) are only relatively slightly broadened when using the Ring-Array, compared to an objective lens system of the same aperture (but having a working distance many times smaller).

In “STED” mode, a torus („donut“)-like intensity distribution around **O** (Fig. 1) is generated in addition to the central focus. In MINFLUX mode (Balzarotti et al. 2017; Gwosch et al. 2020), a donut-like intensity distribution around **O** is generated instead of a central maximum; in “SIMFLUX” mode (Best et al. 2014; Cnossen et al. 2020), patterns generated by structured illumination are used to improve SMLM resolution. The donut half-width is the width (D) of the donut in the inner region. In contrast to [Birk et al., 2017], here the possibility of a STED/MINFLUX compatible donut generation by a flat Ring-Array arrangement is described; by this the large working distances required in a FIB-SEM are realized, while integrating the devices required for FIB-SEM, as well as embedding the specimen in ice. A realization of patterns of structured illumination can be achieved, for example, by interference of a few waves (Heintzmann & Cremer 1999; Gustafsson et al. 2000; Birk et al. 2017).

### Experimental Feasibility of Ring-Array Microscopy

In Supplementary Material, some fundamental challenges shall be addressed concerning the experimental feasibility of optical resolution enhancement by Ring-Array microscopy. In this respect, two major problems have to be overcome: a) The constructive interference of the individual coherent beams emitted from the Ring-Array; b) the collection of a sufficient number of photons at very large working distances. Another experimental challenge (beyond the scope of this general conceptual description) will the appropriate number and distribution of the individual coherent sources in the Ring-Array suitable for a given application. For example, it should be interesting to know whether already a single “ring line” of sources might be sufficient for a given mode of focal/donut intensity distribution, instead of the “homogeneous” distribution of sources in a broad ring zone assumed in the numerical simulations presented here. For example, fiber optics for constructive interference of individual coherent beams have been described (see Supplementary Material), currently allowing phase shift modulation with a resolution down to the femtometer range.

To summarize, compared with the state of the art, the key difference of the Ring-Array-Microscopy concept presented here is the extremely large working distance of a fluorescence-based high-resolution imaging device in an integrated FIB-SEM or another correlative microscope system, providing simultaneously an optical resolution similar to that achievable by objective lens-based devices with low working distance WD and high numerical aperture. Hence the light optical resolution achievable by Ring-Array illumination at extremely large working distances is substantially enhanced compared to imaging with a low NA objective lens. Another major advantage of the Ring-Array method is its flexibility. The use of individual sources according to the Ring-Array principle allows to adjust amplitude, phase, propagation direction, intensity, divergence and polarization individually for each coherent light source. As a result, the focal field distribution can be changed according to specific requirements. For example, in the Ring-Array STED/MINFLUX mode, a first set of sources Si (i = 1,…. N1) may be used to generate a focal diameter for the fluorescence excitation required in STED, while a second set of sources Q_j_ (j = 1,2,… N2) is used to generate a donut intensity distribution around the focal point for STED depletion of fluorescence. In principle, the adjustment of these source parameters may even be used to control the position of the focal spot, implementing beam scanning. Furthermore, compensation of aberrations in imaging systems with high NA (Sheppard and Matthews, 1987) becomes possible, since in principle any apodization function can be synthesized.

## Discussion

Microscopic methods allowing the analysis of large fields of view (FOV) at subcellular resolution are gaining importance in a variety of applications. Typically, they achieve FOVs up to the 1 cm^2^ range and an optical resolution down to about 0.5 µm at a working distance up to several mm (McConnell et al., 2016; Ichimura et al., 2021). The Ring-Array illumination concept presented here is envisaged to be used advantageously wherever still larger working distances combined with a still more enhanced resolution are required: The various illumination modes possible with the Ring-Array arrangement may be applied to measure at extremely large working distances (up to several cm) in correspondingly extensive FOVs the size and shape of optically isolated fluorescent nanostructures (“Ring-Array Nanosizing”); and to realize at selected sites an optical resolution enhancement far beyond the limits of “conventional” low NA/large WD microscopy. Such methods may be applied to flat specimens (e.g. in direct Correlated Light Electron Microscopy/dCLEM); or to large cellular monolayers (or physical tissue sections, or surfaces) in correlative fluorescence microscopy with low NA/large WD objective lenses (and correspondingly extended Fields of View): In a schematic example, such a specimen might be subjected to subsequent modes of imaging: In a first step, a large field of view is screened at micrometer optical resolution using a low NA objective lens, to identify objects of interest (e.g. bacteria, viruses, cells with specific shapes, marker intensities, chromatin patterns etc.). The optical resolution achievable by such a low NA lens screening may be enhanced by a factor of 2, using the SIM mode according to the state of the art. In a step 2 (applicable to optically isolated objects such as viruses scattered on a surface), the Ring-Array-Nanosizing option may be applied to analyse the size and shape of these objects down to the ∼ 100 nm range; in a step 3, selected objects with apparent sizes above this range (e.g. aggregates of 100 nm sized elements) may then be subjected to the focal Ring-Array illumination mode (optical resolution ∼ 200 nm). In additional steps, a super-resolution Ring-Array mode like STED may then be applied to obtain from selected objects an optical resolution down to the tens of nm range. Eventually, even single molecule microscopy related methods (SMLM, MINFLUX, SIMFLUX) might be implemented in suitably advanced Ring-Array Microscopy systems; the required adaptation of the illumination area to selected sites may be realized, for example, in the focusing mode. To create a suitably large illumination area, according to Fig. 4 illumination sources from an inner ring zone may be used; due to the lower synthetic aperture realized in this way, the focal diameter may be increased, e.g. to the few µm range. In this way, the emission characteristics and possibly even the positions of single suitably fluorescent molecules can be determined in specific object regions of interest with small dimensions; thus the nanostructure of such small object regions (e.g. a virus, or a macromolecular complex, or a nuclear gene domain) can be analyzed in more detail. Combining focal illumination with a Ring-Array mode producing a donut or another structured illumination pattern, under optimal optical and labelling conditions the localization precision might theoretically be further enhanced down to the 1 nm limit (Cremer & Birk, 2022).

A great advantage of such Ring-Array Microscopy based procedures would be that the same instrumental device may be used to study a specimen at extremely large working distances at various resolution levels, from the µm to the nm range. Thus it may be regarded as a modern version of the objective lens revolver introduced a century ago into advanced light microscopy systems exactly for the same general purpose: to facilitate and speed up analysis. Such a combination may, for example, quite significantly simplify and accelerate the identification and analysis of individual pathogenic bacteria or viruses (e.g., Cremer et al., 2011; Cremer et al. 2014); or of cancer cells in a tissue section (e.g., Oleksiuk et al. 2015; Lang et al. 2021).

In principle, the large working distance of Ring-Array Microscopy should allow enhanced resolution also of very thick transparent objects, such as extensive tissue volumes, organoids, or spheroids, provided adequate optical conditions (transparency, homogeneity etc.). This will require special additional efforts, e.g. tissue clearing, combination with light sheet microsocopy, axial tomography etc. (Schneckenburger et al., 2020).

Similar advantages as for biomedical applications also arise in material analysis, e.g. contact-free microscopic inspection of surfaces or electronic components: With a large field of view but low numerical aperture, locations of possible defects are positioned, whose nanostructure is analyzed in more detail in additional steps at enhanced resolution using Ring-Array Microscopy. As far as the analysis has to be performed in air or vacuum, novel fluorophores like nanographenes (Liu et al., 2020) may be useful.

## Acknowledgments

We thank Dr. Wladimir Schaufler for valuable discussions and critical reading of the manuscript. The general description of the Ring-Array Microscopy concept presented here is based on the content of a patent application “Verfahren der Ring-Array-Beleuchtungsmikroskopie mit großen Arbeitsabständen und hoher Auflösung” („Methods of Ring-Array illumination microscopy with large working distances and high resolution“), by J. von Hase, B. Humbel, U. Birk, and C. Cremer, filed January 7, 2021 (DE10 2021 000 060.9 2022/January 4, 2022 (PCT WO 2022/148715A1, publ. July 14, 2022 (in German).

