## Supplemental Material for "Ring Array Illumination Microscopy: Combination of Super-Resolution with Large Field of View Imaging and long Working Distances"

### Supplementary Material

#### Units, abbreviations and symbols used

1 nanometer = 1 nm = $10^{-9}$ meter

1 micrometer = 1 µm = $10^{-6}$ meter

NA = Numerical aperture (specification of the objective lens); also used for the corresponding synthetic aperture of a Ring-Array illumination configuration

$\lambda$ = Wavelength of light (vacuum), also instead of the specifications $\lambda_{exc}$ , $\lambda_{MINFLUX}$ $\lambda_{SIMFLUX}$, or $\lambda_{STED}$

$\lambda_{exc}$ = Wavelength of the light used for excitation/illumination (vacuum)

$\lambda_{MINFLUX}$ =Wavelength in MINFLUX Mode

$\lambda_{SIMFLUX}$ = Wavelength in SIMFLUX Mode

$\lambda_{STED}$ = Wavelength in STED mode

G = constant for determining the optical resolving power of objective lenses

n = refractive index of the medium

α: The angles denoted by α are the angles between the Optical Axis and the emissions coming from each source $S_i$ as shown in Fig. 1 Maintext (beam direction defined by the direction of the wave vector), where the vertex of α is the focus center **O** as shown in Fig. 1; $\alpha_{min}$ is the angle between the optical axis and the wave vector of the innermost beam (smallest distance of a source $S_i$ on the Ring-Array according to Fig. 1) from the optical axis; $\alpha_{max}$ is the angle between the optical axis and the wave vector of the outermost beam (largest distance of a source $S_i$ on the Ring-Array) from the optical axis).

a.u. (arbitrary units) = arbitrary units

dCLEM = direct correlative light and electron microscopy

dCLOM = direct correlative light optical microscopy

Depletion = quenching of fluorescence by stimulated emission

EM = Electron Microscope(y)

FIB-SEM = Focused Ion Beam - Scanning Electron Microscope(y)

FWHM = Full-Width at Half-Maximum

MINFLUX (MINimal emission FLUXes) = Minimal emission fluxes: Here: a special method of single molecule microscopy using a torus ("donut") shaped intensity distributi

PSF (Point Spread Function) = Point spread function

Ring-Array = Ring array arrangement: arrangement of sources of coherent radiation in a ring array according to the description (Fig. 4).

ROI(Region of Interest) = area of interest/interesting area

SBF (Serial Block Face) = A method for generating high-resolution three-dimensional images in scanning electron microscopy.

SEM (Scanning Electron Microscope(y)

SIM (Structured Illumination Microscopy) = Microscopy with structured illumination

SIMFLUX (Structured Illumination Microscopy with minimal emission FLUXes) = A special method of single molecule microscopy using an intensity distribution as in structured illumination microscopy or other extended illumination patterns.

SMI = Spatially Modulated Illumination

SMLM (Single Molecule Localization Microscopy) = single molecule based localization microscopy = single molecule microscopy.

SRM (Super Resolution Microscopy) = Super Resolution Light Microscopy

STED (STimulated Emission Depletion) = Stimulated Emission Depletion

STEM (Scanning Transmission Electron Microscope) = Scanning Transmission Electron Microscope(s) (RTEM)

TEM (Transmission Electron Microscopy) = Transmission electron microscopy

Torus ("donut") = "hoop-shaped" object obtained by considering around a ring of radius R all points having a fixed distance r with $r<R$ from a circular line of radius R. Here: Local, torus-like light intensity distribution with a distinct minimum (typically a zero) between two maxima at distance R to realize STED/MINFLUX microscopy

WD (Working Distance) = working distance of the objective lens; in Ring-Array Microscopy the distance $L_{array}$ of the object plane (focus) from the Ring-Array plane according to the description.

Ω (solid angle) = area A of a partial surface A of a spherical surface with radius R, divided by the square of the radius R of the sphere, where the partial surface can be of any outline shape.

**Details of a current CLEM procedure**

In correlative light and electron microscopy, often structured illumination microscopy (SIM) is the means to provide an overview image, and focused ion beam scanning electron microscopy (FIB-SEM) afterwards yields the nanometer-resolution EM images. A 3D map of the whole sample is created and the x, y, z coordinates from the regions of interest are registered by the SIM system located outside the FIB-SEM. The high pressure frozen sample can be directly mounted on the cryo table of the FIB-SEM, and the SIM coordinates can be transferred to the FIB-SEM. The 3D map created with the SIM is matched with the grid image, and then used as an orientation guide in the FIB-SEM work. A lamella is then cut out in the region of interest using the focused ion beam. This lamella can be mounted either with a cold needle (Rubino et al., 2012; Mahamid et al., 2015; Parmenter et al., 2016) in analogy to making lamellae for integrated circuit inspection, or with cold tweezers (Schaffer, et al., 2019) on an electron microscope reticle. The lamella is then re-thinned to a thickness of circa 500 nm and transferred to the cryotransmission microscope. Multiple target structures can be prepared from one cryofixed sample.

A very attractive method is to perform the analysis directly in the FIB-SEM, e.g. using the method of de Winter et al. (2013). STEM tomography at 200 keV in a transmission electron microscope (TEM) opened new perspectives for the analysis of thicker areas (up to 1 µm) of frozen-hydrated cells using cryo-electron tomography (Elbaum et al., 2016; Wolf et al., 2017). These results encouraged the use of STEM tomography in an SEM even at low energies down to 30 keV. Recently published advantages in TEM with low accelerating voltage further support this idea (Banhart, 1999; Linck et al., 2016). If STEM tomography is performed directly in the FIB-SEM, the most critical transfer step from FIB-SEM to cryo-TEM in terms of ice contamination can be bypassed.

To prepare a cryo-lamella for analysis by cryo-tomography, a lamella of about 500 nm thickness and 20 - 50 µm length is prepared out with the ion source at the site of interest. At this time, the site is identified using a cryo-light microscope and the coordinates, usually only 2-dimensional, are transferred to the FIB-SEM. After that, the sample must also be transferred from the cryo-light microscope to the FIB-SEM, and the coordinates correlated with the electron image. This process leaves much to be desired in terms of precision of localization, and sample transfer carries the risk of (partial) thawing and ice contamination.

### Objective lenses for integrated CLEM (dCLEM)

Special objective lenses with an extremely large working distance (WD) of up to several cm have been described (e.g. www.microscopyu.com/microscopy-basics/working-distance-and-parfocal-length). According to equations (1) and (2) in the main text, as an example of an objective lens of such large working distance, a numerical aperture of NA = 0.5 and $\lambda_{exc}$ = 488 nm is assumed; this results in a lateral optical resolution of $d_{min}$ = 600 nm and an axial optical resolution $d_{axia}$l = 2 µm (n = 1; Eq. 2a), i.e., values much worse than the optical resolution that can be achieved using light microscopy techniques for objectives with high NA. Accordingly, using such an axial resolution, a small object structure (ROI) to be examined could be confined to an area of approximately 600 nm laterally and 2 µm axially at half the axial position (50 µm) of a 100 µm thick specimen. To prepare the ROI in such a confined area by FIB would require up to $N_{slice}$ = 2,000 nm/10 nm = 200 cuts for an FIB milling thickness of 10 nm; using SIM would still require half ($N_{slice}$ = (2000 nm/2)/(10nm), or about 100 milling steps. The better the axial resolution, the fewer milling steps are required. The better the lateral resolution, the more precisely the target structure can be identified, and targets of interest selected.

Assuming an objective lens of NA = 0.2 in vacuum with a long working distance (imaging wavelength $\lambda$ = 488 nm and n = 1), relations (1) and (2) would yield a lateral optical resolution of $d_{min}$ = 0.61 $\lambda$/NA = 1.5 µm, and along the optical axis (z) a resolution of $d_{axial} = \lambda/(NA)^2$ = 12.2 µm, resulting in an observation volume $V_{obs}$ (NA = 0.2) = 4/3 π x 0.75 x 0.75 x 6.1 µm$^3$ ≈ 14 µm$^3$. For comparison, a typical small target (ROI) studied with FIB-SEM (dimensions, e.g., 0.2 µm diameter, would have a volume $V_{target}$ = 4/3 π x (0.1 µm)$^3$ = 0.004 µm$^3$), and thus would be smaller by a factor of 14/0.004 = 3,500 than the observation volume using the NA above. However, the larger the observation volume becomes compared to the target volume, the longer the search takes until the target (ROI) can be analyzed with FIB-SEM.

An optical microscopy technique for generating observation volumes in a similarly small range according to the prior art would be STED microscopy. For example, the observation volume now achieved in a commercial STED microscope in biological samples for a high numerical aperture (NA = 1.4) is approximately $V_{obs}$ (STED) = 4/3π × 0.03 × 0.03 × 0.3 µm$^3$ = 0.001 µm$^3$, which is close to the desired observation volume according to the example above.

**General Principle of Ring-Array Illumination**

The rationale (Figure S1) for the principal feasibility of optical resolution down to the nm-range even at very large working distances (e.g. several cm) starts from the fact (Rayleigh 1899; Born & Wolf, 1980) that using an objective lens of a given NA, an „infinite number“ of collimated phase matched light waves of wavelength $\lambda_{exc}$ can be focused to a diameter with an FWHM ~ 0.5 x $\lambda_{exc}$/NA (for more precise relations see maintext). In the case of a typical microscope objective lens, the working distance WD for focusing down to this scale is quite small, however, typically in the sub-millimeter range. Using a finite number of collimated waves (e.g. 7,000) emerging from sites on a spherical calotte (Figure S1a) with an appropriate synthetical aperture (corresponding e.g. to a solid angle of 1.25 π and a half aperture angle $\alpha = 67{,}3°$ ($n = 1.518$), an object plane (x,y) focal diameter around 160 nm ($\lambda_{exc}$ = 488 nm) and a correspondingly small (x,y) donut width (240 nm) may be achieved also at a very large WD, e.g. 4 or 5 cm (numerical simulations, Birk et al., 2017).

If an interior zone of the light sources in the synthetic aperture illumination (Figure S1a) is eliminated (e.g. by assigning in numerical calculations to the sources in this interior zone a zero intensity), close to the nominal focus point, only relatively minor changes in the focal intensity distribution are expected, as compared to the full spherical calotte (Fig. S1b). To test this expectation, numerical simulations were made on the basis of the algorithms used in Birk et al. (2017) under the assumption that the phase matched beams emerged from sites on a ring calotte of spherical shape limited by an internal minimum angle $\alpha_{min}$ and a maximum angle of $\alpha_{max}$ with respect to the optical axis (Fig. S1c). This means that the interior of the ring-like calotte thus produced did not contribute to the illumination. The results obtained by this procedure may be summarized as follows:

1) Using a scalar approximation for the numerical simulations, the axial FWHM(z) produced by such a synthetic calotte illumination scheme with a large but finite number of phase matched beams is approximated with high accuracy by the the well known relation $FWHM_z = (G \cdot \lambda_{exc}) / [n - \sqrt{(n^2 - NA^2)}]$. (eq. 2b, Main Text).For example, with $\alpha_{min} = 45°$, $\alpha_{max} = 70°$, and $n = 1$, an axial $FWHM_z \sim 2.5\ \lambda_{exc}$ is obtained (Table 1 Maintext); this halfwidth is larger by a factor 1.8 as that obtained for a full spherical calotte ($\alpha_{min} = 0°$, $\alpha_{max} = 70°$) with $FWHM_z \sim 1.35\ \lambda_{exc}$. Compared, however, with the axial halfwidth of about 100 $\lambda_{exc}$ obtained with a very large WD objective lens with $n = 1$ and $NA = 0.1$ ($\alpha_{min} = 0°$, $\alpha_{max} = 5.7°$), the calotte derived $FWHM_z$ is smaller by a factor 40.

2) In fully synthetic aperture simulations of intensity distributions along the optical axis (Birk et al., 2017), under certain conditions relatively high side maxima were obtained around the main maximum for long coherence lengths; these side lobes could be substantially diminished, however, by lowering the coherence length appropriately. To which extent such effects occur also in Ring-Array illumination arrangements with a Ring-Array source free interior zone has still to be explored; we assume, however, that using also in this case side lobes

may be diminished in a similar way.

3) For the intensity distribution in the object plane (x,y), a numerical simulation (scalar approximation) assuming n = 1 and about 900 sources distributed in a full ring-calotte with $\alpha_{min}$ = 45° and $\alpha_{max}$ = 70° resulted in a focal intensity distribution of $FWHM_{xy} = 0.45\lambda_{exc}$. Applying the same algorithms for a full calotte with $\alpha_{min}$ = 0° and $\alpha_{max}$ = 70° (about 1,600 sources), a halfwidth of 0.5 $\lambda_{exc}$ was obtained, i.e. practically the same value as established for a glass objective lens of the same NA (NA = 1·sin[70°] = 0.94; compare eq.1 Maintext), with G = 0.47, and in accordance with other simulations performed with slightly different assumptions (Table 1 Main Text).

4) From the numerical results obtained, the volume of the focal intensity distribution (i.e. the observation volume $V_{obs}$) theoretically obtainable at extremely large working distances (e.g. 4 cm) by synthetic ring calotte illumination ($\alpha_{min}$ = 45° and $\alpha_{max}$ = 70°, n = 1) is $V_{obs} = 4/3\ \pi\ (FWHMxy/2)^2 \cdot FWHM_z/2 = 4/3\ \pi \cdot (0.45/2\ \lambda_{exc})^2 \cdot (2.43/2)\ \lambda_{exc} = 0.26\ \lambda_{exc}^3$. This observation volume is somewhat larger (by a factor ~ 2) than the $V_{obs}$ obtained for a full synthetic calotte ($\alpha_{min}$ = 0°, $\alpha_{max}$ = 70°). This figure may be compared with the observation volume (a measure for 3D resolution) of an extremely long WD lens (e.g. NA = 0.1; n = 1) of $V_{obs} \sim 2{,}700\ \lambda_{exc}^3$, which turns out to be larger by a factor of ~ $10^4$ (eq.1 and 2a main text); for a lens with NA = 0.28 (WD = 3.4 cm), the observation volume $V_{obs}$ (NA = 0.28) is smaller by a factor of 60, compared with NA = 0.1.

To summarize, numerical simulations confirmed that using a synthetic ring calotte with a large interior void of sources $S_i$, the focal intensity distribution (and hence the observation volume $V_{obs}$) still closely correspond to that obtained by a full spherical calotte of the same synthetic aperture.

The next step of the rationale (Fig. S1d) is based on the consideration that the focal intensity distribution exclusively depends on the local interference of the waves; hence the same distribution as in Fig. S1c) is expected if the beams are assumed to be emitted not by sources $S_i$ from a spherically shaped calotte but by virtual sources $S_i$* positioned at the intersection points of a plane through the beams, having the same directions and phase relationships as the original beams. If so, the virtual sources in the intersection plane in Fig S1c) may be replaced by real coherent sources of appropriate direction and phase emerging from a flat „Ring Array“.

**Calculation of the intensity distribution generated by the Ring-Array arrangement**

For the practical realization of high-resolution light-optical imaging using Ring-Array arrangements according to the concept described in the main text, it is essential to be able to calculate the intensity distributions generated by them under certain conditions in the object plane and thus optimize their production.

A comprehensive "classical method" for computing intensity distributions in the context of electromagnetic wave theory was described by (Richards and Wolf 1959). Essentially, the solution results from a series of integrals $I_0$, containing products of trigonometric functions, a Bessel function with a product of two trigonometric functions in the argument, and a complex exponential function with a product of two trigonometric functions in the argument. The Bessel functions themselves are not elementary functions, i.e., they must be specified as numerical approximations. The same approach has been successfully applied to calculate the constructive focusing of coherent light in confocal laser scanning fluorescence microscopy as well as in super-resolution confocal laser scanning 4Pi microscopy using two objective lenses with high NA (S. Hell and Stelzer 1992a; S. Hell and Stelzer 1992b) and also for other configurations of polarizations and apertures (e.g. Dorn et al. 2003).

While these integral solutions are very elegant and have been shown to satisfactorily describe focusing by various arrangements of glass lenses, it seems difficult to use them to calculate the intensity distribution produced by a Ring-Array arrangement, which is generated by a finite but typically large number of individual, especially collimated, beams emitted by light sources placed at specific positions, with individually determined individual radiant powers/intensities, phases, polarization and propagation directions. For this reason, a more elementary and flexible prior art calculation method is used here:

Each source $S_i$ (Fig. 1 Main Text) is considered as a starting point of a plane electromagnetic wave, so that the illumination in the object plane is the sum of the interferences of all these plane waves. It is important that the direction of propagation (given by the wave vector) of these waves is the center (**O**) of focal or toroidal (donut) distribution to be generated, and that the phase adjustment of all waves is appropriately made, so that, for example, the phase at the center of focus (**O**) is identical for all waves (apart for a multiple of the wavelength used); further, that a specific state of polarization of the beams is chosen (e.g. linear, radial, azimuthal) depending on the application. The radiant power/illumination intensity of the individual waves can be the same everywhere, or can be adjusted individually. The sum of all waves at a point of interest around **O** is the amplitude and its square corresponds to the intensity around **O**. In calculating the polarization direction of a wave that does not propagate parallel to the optical axis, one proceeds, on the one hand, by rotating a plane wave parallel to the optical axis with a suitable rotation matrix to a wave that propagates perpendicular to the angle of incidence under consideration. On the other hand, the phase difference at a point which is not the focus center, but whose intensity value one is equally interested in, is to be calculated relative to all emanating waves and to be summed up over them.

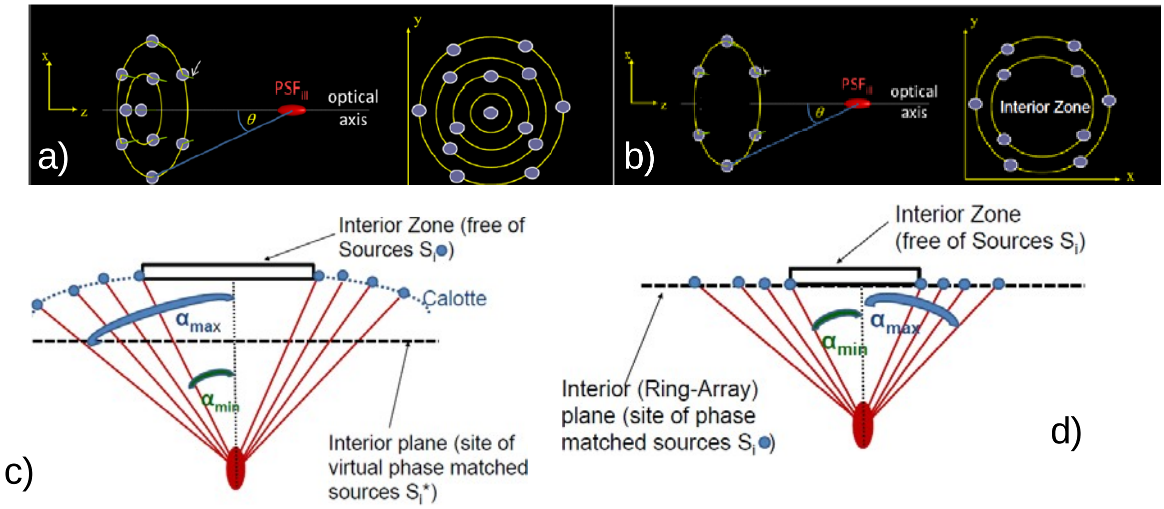


**Fig. S1: Rationale of Synthetic Aperture Ring-Array Illumination**

a) *A spherical calotte with an appropriate synthetical aperture (corresponding e.g. to a solif angle of 1.25 π, a small focal diameter may be achieved also at a very large WD ,e.g. 4 or 5 cm (numerical simulations, from Birk et al., 2017).*

b) *If an interior zone of the light sources in a) is eliminated (e.g. by assigning in the numerical calculations to the sources in this interior zone a zero intensity), only minor changes in the focal intensity distribution are obtained (Fig. a) modified).*

c) *Since the focal intensity distribution exclusively depends on the local interference of the waves, the same distribution as in b) is expected if the beams are assumed to be emitted not by sources $S_i$ from a spherically shaped calotte but by virtual sources $S_i$*(according to the Huygens principle) positioned at the intersection points of a plane through the beams, having the same directions and phase relationships as the original beams.*

d) *The virtual sources in the intersection plane in c) may be replaced by real coherent sources of appropriate direction and phase emerging from a flat „Ring Array“.*

The result of this vector approach can be compared with two limiting cases. On the one hand, for an increasing number of rays, the result is expected to be similar as calculated by [Richards and Wolf, 1959], where the focal field distribution of a theoretical (aberration-free) objective lens is approximated. On the other hand, for a large number of rays distributed over a very small solid angle $\Omega_{array}$, the results are expected to correspond to those obtained for low NA objective lenses. In such a case, vector addition of polarizations is not required, making the problem scalar in nature.

The examples of intensity distributions given in Figs. 5 - 7 and Tables 1 and 2 (Maintext) were calculated in this way according to the state of the art. For more details see [Birk et al., 2017; von Hase et al., 2022].

**Focal intensity distribution by Ring-Array illumination**

Using a suitable distribution of constructively interfering coherent beams produced by the Ring-Array arrangement (e.g. working distance WD 3.4 cm; or 5 cm; or any other distance required to integrate the high-resolution fluorescence mode directly into a FIB-SEM system or into another optical system), a focal intensity distribution is generated that is very similar to that obtained by focusing a laser beam with a Gaussian profile through a lens with a high numerical aperture (e.g., NA = 0.9) at a short working distance (170 µm).

In all examples calculated for focused intensity distributions (Figs. 5, 7, 8; Table 1) there are relatively small differences in the focusing properties of the Ring-Array in vacuum/air; or of focusing in a transparent material (e.g. ice); in this latter case it is assumed that the sample under investigation is embedded in a thin layer of material (n = 1.31) according to the state of the art FIB-SEM techniques. For the Ring-Array concept, it is particularly relevant that also relatively large interior angles (here as an example $\alpha_{min}$ = 45°) and hence correspondingly large Interior Zones are compatible with focusing down to the $\lambda_{exc}$ scale. In principle, with this technique the working distance WD can be chosen to be "arbitrarily" large (e.g. 3, 4 or 5 cm), and a considerable part of the space around the optical axis (Figs. 1-3, 8) can be left out for other components without significantly affecting the focusing. In the example configuration shown in Fig. 3, for WD = 3.4 cm and $\alpha_{min}$ = 45°, a diameter of the

Interior Zone of $D_{interior}$ = 2 x 3.4 x tg[45°] cm = 6.8 cm is obtained.

Figs. (5, 7, S6) and Table 1 show the results of numerical example calculations for the focal intensity distributions generated according to the invention around the focus center **O** (Figs. 1,2) by using a Ring-Array with linearly polarized light sources, while maintaining a large working distance WD and leaving out the inner angular region ($\alpha_{min}$ = 45°, $\Omega_{center}$ = 1.85sr = 0.59 π for other components (e.g., needed for the particle/electron optics of a FIB-SEM, or for other optical systems, such as a low NA objective lens according to Fig. 3). The small differences in the intensity distribution in the object plane (x, y direction) result from the use of a linear polarization of the emitted coherent beams.

Detailed numerical calculations (von Hase et al., 2022) for different thicknesses of the material (ice) surface (up to about 10 $\lambda_{exc}$, corresponding to about 5 µm for e.g. $\lambda_{exc}$ = 488 nm) indicate that, at least up to this depth, the intensity distributions generated by the Ring-Array arrangement (Fig. 4) remain almost identical, with only a shift of the focal center **O** by a few $\lambda_{exc}$. The relative brightness of the intensity maxima also remains almost constant in this region.

It is relevant to the image resolution obtained in the Ring-Array concept that the maximum ratio between the intensity of the main maximum and the secondary maxima does not exceed 0.3. This allows a complete deconvolution of the image data obtained when scanning the object with a focal intensity distribution (Hänninen et al. 1995), in such a way that the optical resolution obtained corresponds approximately to that of the half-width (FWHM) of the central intensity distributions ("central peaks") around **O**. Using confocal detection methods (CLSM; Cremer & Cremer 1978; Sheppard & Wilson 1978; Sheppard et al. 1987, 2007; Brakenhoff et al. 1979, 1985), the optical resolution (for FWHMs see Fig. 6a,b; Table 1) can be improved again by a factor of √(2). A further enhancement of resolution might be obtained by the additional introduction of an „Airy disk“ registration mode (Gardill et al., 2022), or other modes to additionally register fluorescence outside the confocal pinhole region (Cremer & Cremer, 1978).

**Torus ("donut") intensity distribution by Ring-Array illumination**

Figs. (6c-e; S6) and Table 2 show the results of numerical example calculations for donut-shaped intensity distributions generated around **O** (Figs. 1 - 2, 8); here, a Ring-Array arrangement with N = 190 azimuthally polarized light sources was assumed, having at the same time a large working distance WD, and omitting the inner angular region ($\alpha_{min}$ ~ 45°) for other components (e.g. needed for the particle/electron optics of a FIB-SEM, or for a low NA objective lens).

As an essential general result of the example calculations documented in the Figures and Tables concerning the intensity distributions generated by the Ring-Array arrangement, it was found that even at very large working distances (e.g., WD = 3.4 cm, or 5 cm), also donut-shaped intensity distributions may be realized with this method that enable super-resolution light microscopy approaches (STED, MINFLUX) that correspond to the results obtained when using objective lenses of high numerical aperture (up to an effective synthetic aperture of NA = 0.95, n = 1).

When screening an object excited to fluorescence by the illumination conditions of Figs 6c-e; Table 2 (donut intensity distribution in the object plane around **O**, Fig. 1), the illumination conditions may be chosen in such a way that they correspond to the MINFLUX method (Balzerotti et al, 2017; Gwosch et al., 2020); scanning an object excited by the focal illumination conditions of Figs. 5; 6a,b; Table 1, with a first excitation wavelength $\lambda_{fluor}$ and additionally with a wavelength $\lambda_{STED}$ according to the donut intensity distributions of Figs. 6c-e; Table 2 for STED mode, provides illumination conditions corresponding to STED microscopy (Hell & Wichmann, 1994; Hell et al., 1999; Ehrenberg 2014; Sahl & Hell 2019).

Under the conditions of the arrangement, polarization, and phase of the sources in the Ring-Array used in Figs. 6c-e), Table 2, the donut-like intensity distribution occurs only in the object plane (xy), but not in the direction along the optical axis (z). However, this might also be achieved by suitable adjustment of the phases of the sources (Birk et al. 2017; Sahl & Hell 2019) in the Ring-Array; thus, an improvement of the resolution in STED/MINFLUX mode appears to be possible also along the optical axis.

**Ring-array illumination of ice-embedded samples in an integrated FIB-SEM.**

Of essential importance for applications of Ring-Array arrangements is the possibility of their use in the analysis of transparent materials (n >1), e.g. for FIB-SEM investigation of samples embedded in ice under cryogenic conditions. When using the Ring-Array illumination mode, the changes in the refractive index given by the transition from vacuum/air (n = 1) to ice (n = 1.31) must be taken into account. As Figs. 5,6; Tables 1,2 indicate, these effects are small.

**Working Distance (WD)**

While maintaining a high (synthetic) numerical aperture (e.g. NA = 0.95), the Ring-Array arrangement allows to increase the working distance ($L_{array}$, Fig. 1) by several orders of magnitude (e.g. by a factor of 3.4 cm/170 µm = 200x, while maintaining essentially the same focal size (half-widths, Fig. 6; Table 1); thus in a scanning device a similar optical resolution is achieved at WD = 3.4 cm as at a working distance of 170 µm by a high numerical aperture objective lens. The same applies to donut-like intensity distributions realized with the aid of the Ring-Array arrangement, e.g. in STED/MINFLUX (Table 2), or using structured illumination produced by counter propagating laser beams.

When using the Ring-Array concept, despite a large working distance (e.g. WD = 3.4 cm), a very small focal diameter can also be applied to induce photochemical reactions in the region of interest (ROI), or to induce other photophysical processes useful for photoswitching in single molecule localization microscopy(SMLM). In addition, such a small focal diameter in combination with suitable illumination intensities may also be used for localized material processing, e.g., laser-induced ablation of small surface irregularities in optical elements.

For example, with a power P = sum (Pi) of the power of the individual beams $S_i$ emitted by the Ring-Array sources in the focal area around **O** (Fig. 1), the average intensity is $I_{average}$ = sum (PI )/$A_{Focus}$ , where $A_{Focus}$ ~ $FWHM^2$ ~ $(d_{min})^2$ . For example, for a working distance of several cm, using a lens with a NA = 0.06 ($\lambda_{exc}$ = 488 nm), Eq. (1) gives a focal diameter of approx. 4 µm ($A_{spotconv}$ = 16 $\mu m^2$); using a focusing Ring-Array arrangement, even at a working distance of e.g. 3 or 5 cm (theoretically even more), a focal diameter about 15 x smaller (0.25 µm) and thus a focal area of about 270x smaller ($A_{Array}$ = 0.06 $\mu m^2$) may be generated. The focal intensity produced by the

Ring-Array illumination would thus be ~ 270x higher at the same total beam power by a factor of $A_{spotconv}/A_{Array}$; with an objective lens of NA = 0.06 in an integrated FIB-SEM or an integrated dCOLM system (Fig. 3), an approx. 270x higher power would thus be required to achieve the same focal illumination intensity at the required large working distances; instead of a laser output power of e.g. 100 mW, one of 27 W would be required which in most cases may be inconvenient.

### Technical configuration of the Ring-Array illumination mode

According to the Ring-Array concept (see Figs. 1, 2), the beams $S_1$ , $S_2$ ,... $S_N$ are positioned in a solid angle $\Omega_{array}$ characterized by the limiting beams $\alpha_{min}$ and $\alpha_{max}$. In contrast to the arrangement revealed in [Birk et al. 2017], $\Omega_{array}$ is limited by the inner zone, which is free of emitting rays (diameter $D_{interior}$); The solid angle of this inner zone (which includes the space between the focal position **O** and the inner edges of the ring array) is referred to as the $\Omega_{center}$ (see Fig. 2). The remaining space free of the beams $S_1$ , $S_2$ ,... $S_N$ is given by $\Omega_{bottom} = 4\pi - \Omega_{array} - \Omega_{center}$ (see Fig. 2; Fig. 8). The illumination of the Ring-Array is performed by appropriately configured coherent beams, which typically produce homogeneous illumination or structured illumination or other adequate intensity distributions to suitably irradiate the positions of the $S_i$ sites; in the following, the $S_i$ positions from which the coherent (e.g., collimated) beams emanate due to the illumination of the ring by the same coherent source (typically a laser) will be referred to as light sources (or sources). In order to generate the required changes in direction, phase, polarization, and intensity of the sources Si, various methods may be selected. From the point of view of simplicity, an appropriately designed diffractive optical element (DOE) or spatial light modulator (SLM) may be used; if necessary, the beam sources $S_i$ may additionally include optical elements to be able to individually change the direction, phase, polarization or power of the emitted beams. Diffractive optical elements (DOEs) have the advantage of allowing a stable and economical realization of a Ring-Array device; the disadvantage is that the ring array may have to be exchanged in order to realize other intensity distributions in the object plane around **O**. However, with suitably large working distances WD = $L_{array}$ (Fig. 1), it should be possible to realize a Ring-Array with multiple sets of sources, e.g., an array set A of sources $S_{11}$, $S_{12}$,…(“structured“), $S_{21}$, $S_{22}$,…(“focal“) for focal intensity distributions for illumination with an excitation wavelength $\lambda_{exc}$; and an array set B with sources $S_{31}$, $S_{32}$,… (“donut“), for generating a donut-shaped intensity distribution for illumination with a wavelength $\lambda_{STED}$.

Spatial Light Modulators" (SLMs) on the other hand have the great advantage of flexibility. For example, they allow rapid adaptation to realize different intensity distributions around **O**. However, the technical integration may be more expensive and more susceptible to mechanical stress. All those implementations fulfill the purpose of the Ring-Array if they produce a desired intensity distribution in an integrated FIB-SEM or another optical device at a given large working distance.

The numerical examples given (Fig.6; Tables 1,2) are based on a Ring-Array arrangement assuming a total number of N = 190 (in Fig. 7b: N = 760) collimated beams, each of suitable phase, angle and polarization, with the intensity of all individual beams assumed to be equal in the examples shown here. Any other annular distribution or number may be used as long as the goal of the Ring-Array concept is maintained to realize a fluorescence illumination system with a solid angle $\Omega_{array}$ using coherent waves that corresponds to a large synthetic aperture relative to the NA of a lens-based microscopy system, but with the essential distinction of a suitably large working distance (WD) and a suitably large central aperture with a solid angle $\Omega_{center}$, in such a way that the devices can be placed in a FIB-SEM or into other microscope systems for undisturbed illumination/detection. In this case, it may be necessary to adjust the phase, direction, polarization, divergence, and power of the beams accordingly in an individual manner. To determine the required parameters, numerical simulations according to the state of the art (see above) may be used.

According to the Ring-Array concept, several implementations of the polarization of sources $S_i$ may be considered. For example, all coherent beams emitted from sources $S_1$, $S_2$ ,... $S_N$ in the Ring-Array (Fig. 4) may have the same polarization direction; in this case, implementation of the polarization conditions would require only a corresponding polarization of the coherent radiation (wavelength $\lambda_{exc}$, $\lambda_{STED}$ used to illuminate the Ring-Array. The polarization conditions, however, can be implemented also by site-specific modification in the Ring-Array (e.g., by appropriately designed diffractive elements or by prior art spatial light modulation).

The special polarization modes (linear, circular, radial or azimuthal polarization of the waves emanating from the sources $S_i$ (i = 1,2,...N) may be realized either in separate designs in each case, or in combinations of suitable sets of sources.

For example, it would also be possible to place in a single ring array area 2, 3, or 4 sets of sources, where, for example, set 1 includes $N_1$ sources with linear polarization; set 2 includes $N_2$ sources with circular polarization; set 3 includes $N_3$ sources with radial polarization; and set 4 includes $N_4$ sources with azimuthal polarization.

The different irradiation modes can be used together (e.g.. $\lambda_{exc1}$ for focusing/CLSM/SMLM; $\lambda_{exc2}$ for STED/MINFLUX), but can also be used individually from each other by means of different polarizations/filters; a further variant is a temporally and/or spatially variable illumination of the ring array with different wavelengths of coherent radiation; in this way, the intensity distribution in the object plane can be further modified via the irradiation modes given as examples (focal, donut).

In the illumination mode examples presented in the following, it is assumed that the radiation emitted by the individual sources $S_i$ of the Ring-Array arrangement, when applying a specific irradiation mode (e.g., focusing, donut; linear/azimuthal polarization), all have the same power (W/s) [$P(S_{11}) = P(S_{22}) = \ldots P(S_{NN})$], respectively, and only the phase/direction is suitably modified. This can be realized in various ways (Birk et al., 2017), including nanolithography; or multiple glass fiber arrangements.

For example, nanolithographically generated Ring-Array arrangements might be constructed to be used in structured mode (e.g. nanoshape analysis); focused mode (e.g. CLSM), Single Molecule Localization mode (creation of a small illumination field at selected sites); or donut mode (e.g. STED/MINFLUX), or in a combination of the different illumination options in the same Ring-Array.

In order to additionally control the power $P_i$ of individual sources or individual sets of sources suitably also independently, e.g., micromirror systems or other elements can be used to control the power of multiple beams.

Moreover, by suitably reducing the number of sources $S_i$, instead of predominantly concentrating the total power emitted by the Ring-Array on a single focal or donut volume, it is possible to generate periodic patterns of such intensity distributions in the object plane, with a spacing greater than the wavelength used for illumination (Figs. 7). This makes it possible to use such periodic intensity distributions advantageously in object screening as well.

**Size of Ring-Array**

For example, the area of a Ring-Array with of $\alpha_{min}$ = 45°, $\alpha_{max}$ = 70° (see Fig. 1), and a working distance WD = 5 cm would be sufficiently large to accommodate e.g. 4 sets of sources with 190 emitters each (e.g. set 1 for structured illumination microscopy; set 2 for focused intensity distribution in confocal laser scanning microscopy; set 3 for

MINFLUX/STED in xy; set 4 for MINFLUX/STED in z-direction): The above configuration results in an outer ring array radius of $r_{exterior}$ = 5 cm x tan(70°) = 13.7 cm, and an inner radius $r_{interior}$ = 5 cm x tan(45°) = 5 cm, i.e., a total area of 511 $cm^2$ = 51,100 $mm^2$. Thus, for each source (total $N_1 + N_2 + N_3 + N_4$ = 4 x 190), there would be 51,100 $mm^2$/(4 x 190) = 67 $mm^2$. To realize such an area size/individual source should be technically feasible with suitable miniaturization (e.g. by using glassfiber optics, see below).

**Nanosizing by structured Illumination**

In many application cases, it may be sufficient to obtain more information about the size and shape of optically isolated small objects (i.e. objects with a next neighbor distance larger than the optical resolution of the imaging system). In these cases, it is not necessary to create focal or donut intensity distributions; hence, the constructive phase adjustment of multiple Ring-Array beams (see below) can be avoided.

For example, both virtual SMI microscopy calculations with counter propagating waves (Failla et al., 2002b) and experimental SMI measurements (Failla et al., 2002a) indicated that using an exciting wavelength of 647 nm ($\theta_{max}$ = 0°), sizes of small particles may be determined with good precision down to diameters of about 40 nm. In these numerical SMI simulations, a maximum wavelength $\lambda_{exc}$ = 647 nm (NA = 1.4; n = 1.515) corresponds to an enveloping PSF with a halfwidth of $FWHM_z = n\lambda_{exc}/NA^2$ = 500 nm, and a fringe distance $\Delta_{min}(z) = \lambda_{exc}/[2n\cdot\cos(\theta_{max})]$ = 647 nm/[2 · 1.515·cos(0)] = 214 nm. Since in the theory of nanosizing (Failla et al., 2002b), only the stepwise movement of a fluorescent nanostructure through a Standing Wave Field and its convolution with a PSF having a halfwidth several times larger than the fringe distance is assumed, the virtual microscopy results may be used also in the case a) that the fringes are generated not along the optical axis but along the object plane; b) that instead of moving the nanostructure stepwise along the optical axis, it is moved in the object plane orthogonally to the standing wave field; and c) that the FWHM to be used for object plane nanosizing is not the $FWHM_z$ along the optical axis but the $FWHM_{xy}$ along the object plane. Hence, the virtual microscopy results cited above for axial nanosizing ($FWHM_z$ = 500 nm; $\Delta_{min}(z)$ = 214 nm) should be valid also for a Ring-Array nanosizing mode with a corresponding $FWHM_{xy}$ and fringe distance $\Delta_{min}(xy)$. Such a object plane fringe distance ($\Delta_{min}(xy) = \lambda_{exc}/[2n\cos(90°-\theta_{max}]$ (= $\lambda_{exc}/[2n\sin(\theta_{max}])$) around 200 nm may be generated by Ring-Array illumination using e.g. n = 1.3 (water), $\lambda_{exc}$ = 488 nm, and $\theta_{max}$ = 70°.

In the object plane, a $FWHM_{xy}$ = 500 ($\lambda_{exc}$= 647 nm) corresponds to a Numerical Aperture of NA = 0.51 $\lambda_{exc}/FWHM_{xy}$ = 0.65, or to a working distance around 1 cm (https://www.newport.com/f/long-working-distance-objectives).

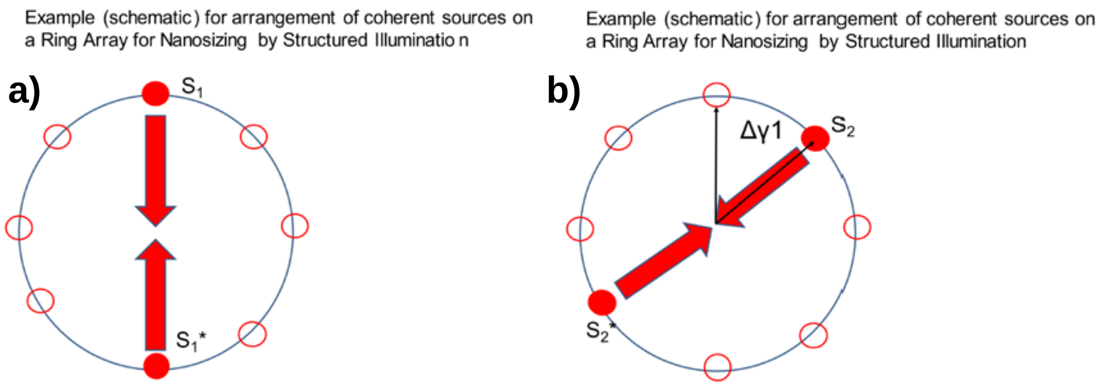


**Fig. S2: Illumination Ring-Array Option for Nanosizing**

a) *Instead of the constructive interference of multiple phase matched beams required for the generation of a focal/donut intensity distribution in the object plane, a single pair of counter propagating coherent, collimated waves from sources $S_1$, $S_2$ is sufficient to create a structured illumination ("standing wave") pattern in the object plane useful for the determination of the size of fluorescence labeled nanostructures.*

b) *Nanosizing and shape determination may be substantially facilitated by rotating the direction of the single beam pairs ($S_2$, $S_2$*) in the Ring-Array by defined angles Δγ.*

Such a "Nanosizing" approach might therefore be used also in Ring-Array microscopy to enhance the size and shape resolution. In contrast to earlier implementations, the fringe distance of the structured illumination pattern is determined by the synthetic aperture of the Ring-Array and thus independent from the NA of the objective lens used for fluorescence detection. Let us assume (Fig. S2a) that in the Ring-Array, coherent collimated beams are emitted only by two opposed sources $S_1$, $S_1$*. By interference, this results in a standing wave field (indicated by the two counter opposed arrows) and may excite the fluorescence on an optically isolated structure of diameter s in the object plane.

Using high NA objective lenses, the creation of harmonic illumination patterns in the object plane by the imaging of two coherent beams is an experimentally well established method (e.g. Heintzmann & Cremer, 1999; Gustafsson, 2000). Recently, harmonic illumination patterns have also been used for Retina diagnostics in structured illumination ophthalmology (Schock et al., 2022), i.e. under the low NA conditions of the human eye.

However, in this way sufficiently small fringe distances $\Delta_{min}$(xy) for nanosizing may be obtained only with high NA objective lenses and hence small WDs. The Ring-Array illumination approach allows to separate the generation of the patterned illumination from the objective lens used for fluorescence detection.

By scanning e.g. the stage with a given step size and registration of the images obtained (e.g. by a low NA lens), an image may be obtained similar to Spatially Modulated Illumination (SMI) microscopy (Hausmann et al., 1997; Failla et al. 2002a,b, 2003; Albrecht et al., 2002; Baddeley et al., 2007; Cremer & Birk, 2022). Under the conditions of Ring-Array illumination, smallest fringe distances ($\lambda_{exc}$ = 488 nm n = 1.3, $\theta_{max}$ = 70°) of the interference maxima along the object plane of $\Delta_{min}(xy) = \lambda_{exc}/(2n\sin\theta_{max}])$ = 200 nm would be obtained.

Using the rationale for virtual SMI nanosizing outlined above, numerical simulations have been made for Ring-Array illumination based nanosizing, assuming a Ring-Array aperture angle (Fig. S2) of 70°, and objective lenses for fluorescence detection with a) an NA = 0.35 (WD = 1.3 cm) and b) with NA = 0.28 (WD = 3.4 cm), respectively. For an excitation wavelength of 647 nm and n = 1 (air/vacuum), or n = 1.3 (water), respectively, an optimal size resolution between 65 nm and 80 nm was estimated (see below, Fig. S3).

In a further implementation (Fig. S2b), instead of the sources $S_1$, $S_1$*, the sources $S_2$, $S_2$* positioned at an angle $\Delta\gamma_1$ with respect to $S_1$, $S_1$* may be „activated“ to emit collimated beams. The process of step 1 is repeated and provides nanosize information in the new direction given by the angle $\Delta\gamma_1$. By repeating the procedure, a shape profile of the object is obtained, similar to the principle of computer tomography.

In this approach, the phase difference between the individual pairs of counter propagating beams is not important for the nanosizing process; for each measurement, the phase difference between the two beams has just to be stable during the scanning process (either stage scanning; or corresponding conditions for phase scanning, as in Lemmer et al., 2008). Since the ”switching ON“ of the sources $S_i$, $S_i$* can be done very fast (e.g. in fiber optics based illumination), shape profiles of optically isolated nanostructures may be acquired in a short time. For example, if one assumes a stage scanning with step sizes of 20 nm, a total of 50 steps and 0.1 s illumination time for each step (Reymann et al., 2008), a few seconds would be sufficient for a size measurement in one specific direction, and 1 min for a shape measurement. Such nanosizing/nanoshape measurements may be automated and would be feasible simultaneously for all nanostructures of the same type in the field of view of the low NA lens used for fluorescence registration (see below for an application perspective in genome biology).

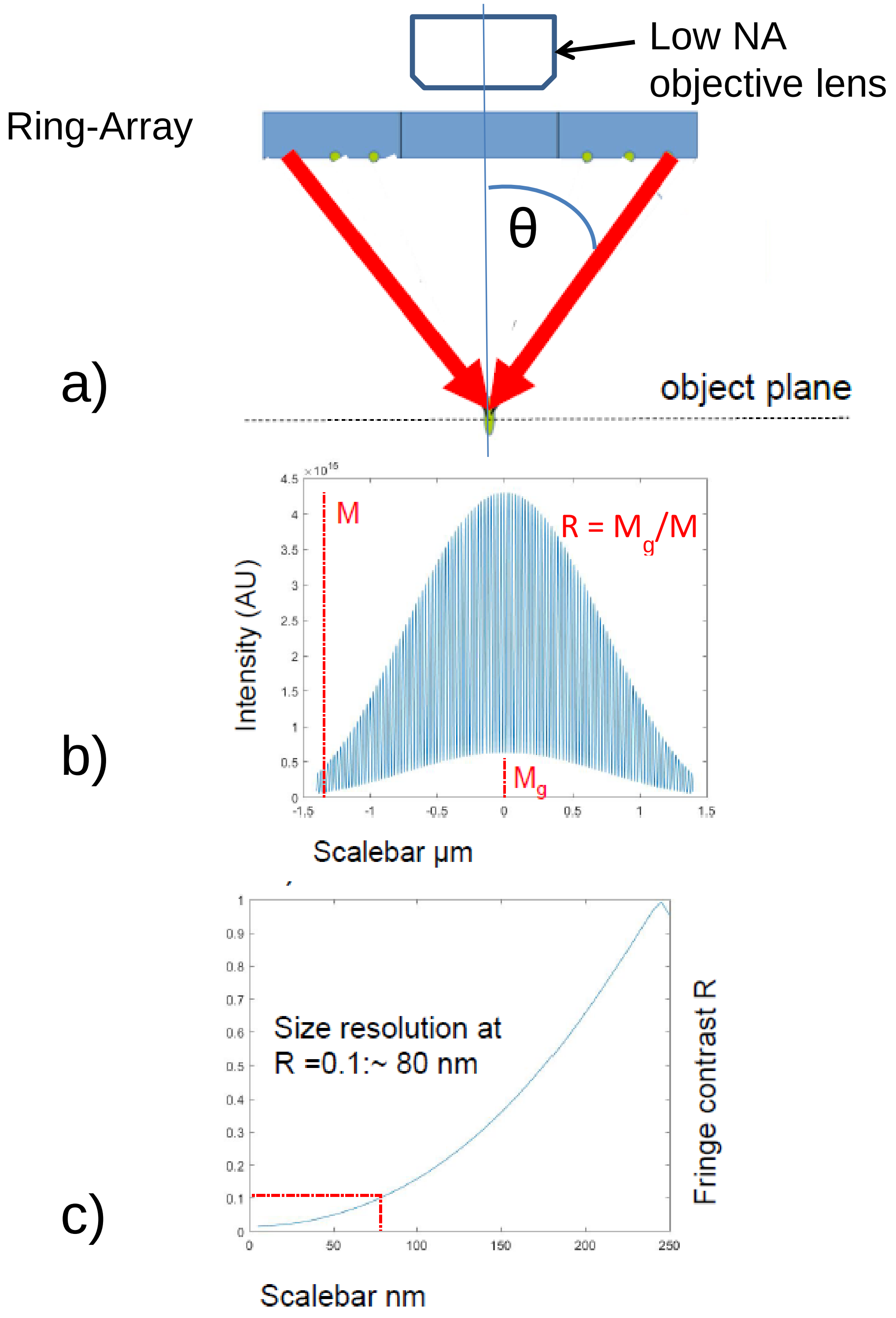


**Fig. S3: Ring-Array-Nanosizing at extremely large Working Distances**

a) *Instead of Ring-Array illumination with multiple phase matched beams to generate e.g. a focal or donut like intensity distribution in the object plane (Fig. S1d), at a given time only a pair of two counter propagating coherent beams emerges from two opposed sites of the Ring-Array (Fig. S2). The two beams (angle θ between the beam direction and the optical* *axis e.g. 70°) create a standing wave field (harmonic intensity distribution) in the object plane (coordinates x,y), with a fringe distance $d_{xy}$ several times smaller than the lateral resolution ($FWHM_{xy}$) of the low NA/large WD objective lens used for fluorescence registration (e.g. NA = 0.28, WD = 3.4 cm), positioned in the interior zone of the Ring-Array. The optically isolated nanostructure (diameter e.g. 100 nm) is moved in defined steps $\Delta_{xy}$ through the harmonic pattern generated in the object plane; at each step the fluorescence image is registered by the low NA lens.*

b) *Expected result (schematic) from measured intensity distribution (numerical simulations) through the center of the fluorescence image of a nanostructure with a diameter (size) of 100 nm, assuming a low NA lens (NA = 0.28), and an angle of θ = 70° (corresponding to a synthetic aperture of the Ring-Array of sinθ = 0.94). Shown is the enveloping PSF of the low NA lens (blue) and the "background" (white) generated by the convolution of the low NA lens PSF with the fringe pattern. From the maximum (M) of the intensity distribution and the "background" (height $M_g$), the fringe contrast $R = M_g/M$ is determined. Scale bar: µm.*

c) *Numerical simulation of the dependence of the fringe contrast R (ordinate) from the size of the object (abscissa; diameter perpendicular to the extension of the fringes in the object plane). In this example, an exciting wavelength of 647 nm and NA = 0.28 (n = 1/air vacuum) was assumed. From this (R = 0.1), a size resolution limit of about 80 nm is obtained. Using the same condition but assuming a refraction index of 1.3 (water), the fringe distance becomes somewhat smaller, resulting in a size resolution of about 65 nm. Instead of stepwise movement of the object, the relative phase difference of the two beams may be stepwise modified. In case of a non spherical object, the shape of the object may be determined by rotating the object and measuring the resulting size in the respective direction of stepwise movement along the object plane. Such a "nanoshaping" may be facilitated considerably by sequential nanosizing with pairs of beams (Fig. S2).Scale bar: nm.*

We conclude that object plane related nanosizing/nanoshaping down to the ≤100 nm range should become feasible using Ring-Array generated structured illumination patterns even at extremely large working distances.

The nanosizing potential of a Ring-Array-SIM illumination using a low NA large WD lens may be compared with the size resolution obtained under the same conditions as for the example mentioned above: With homogeneous widefield illumination, the size resolution (smallest size to be correctly determined) of a fluorescence labelled optically isolated nanostructure in the object plane would be similar to the Airy Disc extension (Baddeley et al., 2010); assuming e.g. an NA = 0.5 and $\lambda_{exc}$ = 488 nm, this would amount to a size of about 500 nm; for NA = 0.28, it would be ~ 900 nm.

To summarize, by enlarging the aperture angle of two beams interfering in the SMI mode by Ring-Array illumination allows to substantially enhance the nanosize resolution of a low NA objective lens. For NA = 0.28, for example, the enhancement may be up to tenfold.

Combined with automated image registration and evaluation procedures, Ring-Array based nanosizing is envisaged to allow size and shape analysis of optically isolated small targets in extremely large fields of view. For example, using a low NA lens (NA = 0.28) with an extremely large field of view of about $1cm^2$ (Tsai et al., 2015), using fast automated procedures, nanosizing of the transcriptional status of appropriately labelled specific gene domains (Cremer T et al., 2015, 2020; Gelleri et al., 2023; Maeshima et al. 2023; Miron et al., 2020), might be performed simultaneously for a hundred thousand individual cells in a few minutes, with total fluorescence registration times of a few seconds (Reymann et al., 2008) for the entire Field of View. In a biomedical application, this should e.g. allow in cellular pathology to register transcription related data of 10,000 specimens (3 mm diameter, with 10 serial sections each placed adjacent to each other) in a few hours ($10^4$ FOVs of 1 $cm^2$ each, fluorescence registration time for each 1 $cm^2$ FOV e.g. 2s). If instead the nanosizing would be performed with a high NA objective lens (structured illumination generation through the lens) and correspondingly small fields of view (typically 0.5 mm Ø), then the scanning time (total area of the 10,000 specimens to be analysed ~ $1m^2$, or 4 x $10^6$ FOVs to be sequentially scanned, each small FOV 2 s) would amount to about 7 weeks registration time.

High Throughput imaging of extremely large surfaces (e.g. 1 $m^2$ with 10,000 FOVs of 1 $cm^2$ each), containing small subwavelength sized fluorescent objects to be measured might be interesting not only for cellular pathology and other biomedical applications (e.g. testing of drugs), but also for applications in material sciences, e.g. to identify small surface contaminations). A laser power in the range of a few hundred mW should be sufficient to excite for this in biological specimen a sufficient fluorescence signal, simultaneously illuminating the entire field of view (Schock et al., 2022): Depending on the specific label chosen (e.g. covering the compact domain core only; or the entire domain), transcriptionally active gene domains are expected to be indicated either by a reduction in core size, or by an increase in the size of the entire domain compared to the silenced domain (Fig. S4). To allow the required optical isolation of the labelled small domains, the resolution of ~ 1 µm achievable with a NA = 0.28 lens would be sufficient to discriminate in the large majority of cases in the cells two domains labelled with the same spectral signature in a nuclear area of 100 $µm^2$.

In many cases, it may be desirable to perform such a high throughput nanosizing not only for one specific type of gene domain but simultaneously for multiple ones. This may be achieved by appropriate “spectral signatures” (Cremer et al., 1999). Presently, 7 types of labelling gene domains can be discriminated by appropriate excitation wavelengths and fluorescence emission filters; combinatorial labelling of 7 spectral signatures allow to identify up to $2^7 - 1 = 127$ optically isolated gene domains (Bolzer et al., 2005).

In addition, it may be noted that low photon yields only are required for SMI-nanosizing; according to numerical simulations (Cremer et al., 2003), a total of about 1,000 registered photons should be sufficient to allow effective nanosizing below 100 nm. This makes possible illumination conditions even suitable for high throughput live cell nanosizing, e.g. of the transcriptional status of specific chromatin domains (Reymann et al., 2008).

Ring-Array based structured illumination of extremely large FOVs are envisaged to be

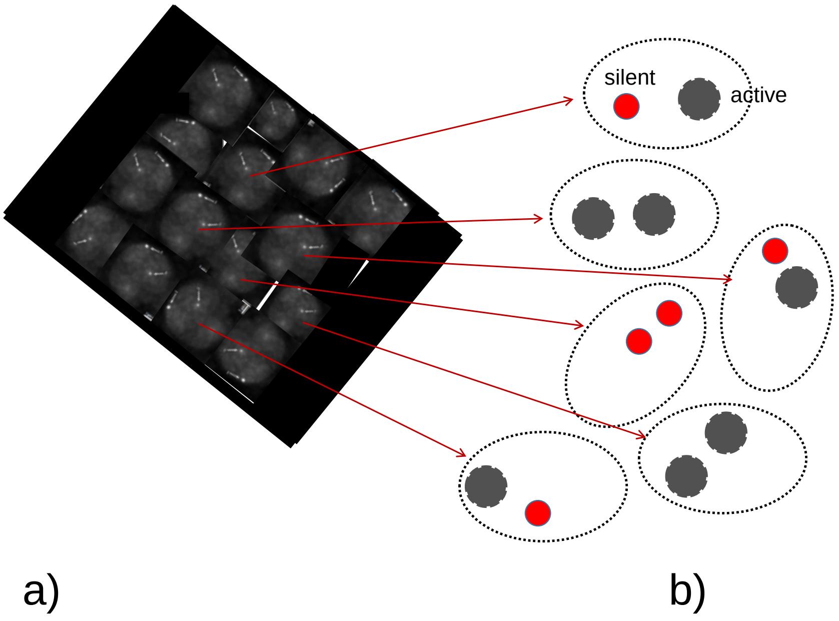


**Fig. S4: Schematic example for high-throughput Nanosizing of transcriptional heterogeneity of specific gene domains in large multicellular aggregates**

*a): Large field of view with high numbers of cells containing fluorescence labeled gene domains (example cells from high NA SMI-Nanosizing, Mathee et al. 2006)*

*b) Right: Result of Ring-Array Nanosizing (assuming e.g. an objective lens with NA = 0.28, $FWHM_{xy}$ ~ 1 µm, WD = 3.4 cm, and FOV ~ 1 $cm^2$). Here, the label is assumed to be performed in such a way that silent gene domains (indicated Red) have a somewhat smaller size (e.g. 70 nm) than transcriptionally active ones (e.g. 100 nm, indicated Gray).*

It is important to state that in contrast to Ring-Array Microscopy with multiple phase matched illumination beams (Figs. 5-7, Tables 1,2 Maintext), the Ring-Array **nanosizing** mode does not provide enhancement of optical resolution in the sense of eq. (1,2). For this, it would be required to sum up the phase matched amplitudes of the various coherent beams emitted from the Ring-Array, as assumed in the general concept. But what is possible in the nanosizing mode, are improved size determinations of optically isolated nanostructures using the SMI principle.

**Phase Calibration for focal/donut Intensity Distributions**

The general Ring-Array illumination concept is based on the assumption that the phases of the coherent beams emanating from the Sources $S_i$ in the Ring-Array can be calibrated in such a way that the required constructive interference is obtained (e.g. a focal intensity distribution, or a donut distribution). This typically needs a limitation of the phase error to values ≤ 2π/10, or (for n = 1) optical length differences ≤ $\lambda_{exc}/10$ between the multiple beams, i.e. in the range of 50 nm (results of numerical simulations, data not shown). At first sight, this requirement appears to be extremely difficult to fullfill. However, due to new technology, it appears that various alternatives exist to meet this challenge.

The generation of multiple coherent beams, in particular also collimated beams with controlled intensity, phase and polarization relationships, is possible in various ways by generating light from a single laser light source, e.g., the individual coherent beams by means of a microlens array and guiding them through suitable mirror configurations to the desired positions of the point sources $S_1$, $S_2$ ,...$S_N$ in the Ring-Array arrangement (Fig. 4). This can be e.g. realized a) using free-space optics, or b) using a fiber optics based approach. Such a microlens configuration can be used together with a liquid crystal array in transmission mode to allow control of the phase relationship at the location of the point sources $S_1$ , $S_2$ ,...$S_N$. By introducing intensity-regulating elements (e.g., gray filters, polarizing elements, acousto-optical modulators, etc.), it is additionally possible to control also the intensity of the beams individually; in this way, in addition to the intensity distributions (focusing, donut) given in the examples presented (Figs. 5-7; S6; Tables 1,2), other microscopy modes can be realized using the same Ring-Array arrangement according to the invention, such as light sheet microscopy, optical projection tomography or axial microtomography.

Another especially attractive solution to the phase matching problem is the use of optical fiber arrangements to generate multiple coherent partial beams with individually adjustable phase relationships, intensities and polarizations (Birk et al., 2017). This can be realized relatively economically with suitably programmable fiber optic arrays. Such fiber arrays ["Fiber optical singlemode switches/splits (polarization maintaining)"] are presently commercially available with up to 16 outputs, but this number can be easily increased. The phase relationships of the coherent light beams emitted by the fiber optic bundle can be programmed/implemented in a desired manner using the programming/calibration technology provided. To fully realize the focal/donut intensity distributions simulated (Figs. 5 - 7; Table 1,2), less than 200 separate beams would be required.

In principle, all manipulations of laser beams required for Ring-Array illumination- based correlative microscopy, such as beam splitting, polarization control, optical switching, phase control, etc., can be performed with the above fiber optic arrangements. Fiber optic technologies have unique advantages: they are environmentally stable (e.g., the required interference beam patterns can be kept very constant), flexible, robust, and economical. Therefore, all-fiber beam realization is one of the optimal options for implementing the constructive interference of the Ring-Array beams required for optical resolution enhancement. For this, a variety of laser sources can be considered as light sources for fiber coupling according to the Ring-Array concept, ranging from pulsed light sources to CW lasers in the entire visible and near UV range. The polarization state of the beam can also be preserved by using polarization preserving fibers [e.g., polarization preserving single mode optical fiber (www.thorlabs.com)]. For example, to generate the coherent light sources $S_1$, $S_2$,...$S_N$ of the Ring-Array illumination from a given laser source, cascaded optical fiber beam splitters may be used (e.g., https://www.china-tscom.com/products/plc-splitter). Phase control can also be performed according to the state of the art, e.g., using a fiber optic phase shifter (www.phoenixphotonics.com).

The use of multiple collimated beams in the Ring-Array arrangement has several advantages: a) the use of collimated beams can in principle be realized by an arbitrary spacing of the light sources $S_i$ emitting the collimated beams, by using appropriately dimensioned Ring-Arrays; b) with increasing distance from the origin, the number of sources $S_i$ of the collimated beams can be made very large; c) the number, intensity and spatial distribution of the light sources $S_i$ (using, for example, microlenses, or Spatial Light Modulators) can be adjusted individually. For example, by adding a possibility to control the intensity $I_i$ of the collimated beam emitted by a light source $S_i$ (e.g. in a way similar to a projection beamer), the light distribution can be individually modified in a fast and efficient way.

To obtain constructive interference of the beams emitted by the Ring-Array sources, the phases have to be adjusted accordingly in such a way that, for example, when focusing, the intensity at a point **O**, which will be the center of focus, becomes a maximum with linear polarization settings. For such a distribution, this may be implemented as follows:

The coherent beams emitted by the N light sources $S_1$, $S_2$ , ...$S_N$ (e.g., N = 190) are divided into N/2 pairs of counter-propagating beams (e.g., N/2 = 190/2 = 95). For example, in Fig. $S_1$, two such pairs S1, S1* and S2, S2* may be considered. As a first step, the relative phase between the beams from $S_1$ and $S_1$* is adjusted in such a way that the intensity of the detected signal obtained by the detection optics using a subwavelength-size calibration object (e.g., a fluorescent "nanobead" with a diameter of 100 nm or smaller) at a given position in the object plane (e.g. **O**, Figs. 1-3,8) is maximized (Failla et al. 2002; Baddeley et al. 2007; Lemmer et al. 2008; Reymann et al. 2008). At step 2, this procedure is repeated with another pair $S_2$ and $S_2$* of counter propagating beams (after switching off the intensity of the beams of step1). The calibration steps are repeated for the counter propagating waves of all 95 pairs of the 190 sources. At sufficiently stable maintenance of the individual calibration made, simultaneous switching ON the intensity of all the 95 calibrated beam pairs, a focal intensity distribution as obtained e.g. in Fig. 5; 6a,b; Table 1) is expected.

If one then wants to obtain a donut shaped intensity distribution to realize STED/MINFLUX microscopy (Fig. 6c-e; Fig. S6; Table 2), for example, one only needs to adjust the polarizations of the waves emitted by the sources of the Ring-Array azimuthally. Other implementations according to the Ring-Array concept by appropriate parameter selection are also possible. To facilitate this process, the alignment of the phases may be performed in pairs with waves propagating in opposite directions.

Hence, in the callibration example given, the phase adjustments are performed sequentially one after the other. If necessary, additional adaptive optics devices according to the state of the art (e.g. www.thorlabs.com/Adaptive_Optics) may be used.

Since the process can be automated, the calibration can be performed in a short time. For example, using 100-nm beads ("nanobeads") with good fluorescence emission, a sufficient photon yield can be obtained in 100 ms; if 30 measurements are taken at a time to optimize the phase of a given pair of counter-propagating waves, a total of e.g. 30 x 95 x 0.1 s ~ 300s is required, assuming a sufficiently stable mechanical stage (e.g., www.pi-usa.us).

In objects with refractive index variations, adaptive optics techniques can be used to perform phase optimization. This is the case, for example, when focusing is performed with variable ice layer thickness.

**Sample mounting and detection of the fluorescence signal**

There are various options for the spatial arrangement of the sample holder and optical components used to detect fluorescence excited in and emitted from the sample by means of the Ring-Array illumination (for a general schematic example, see Fig. 8). A simple but effective method for detecting fluorescent light is to use an objective lens with a large working distance. Since the focal volume in which fluorescence is generated is defined by the spatial distribution of the illumination intensity, when scanning the object point-by-point, the critical variable for the detection light path is the total amount of fluorescence signal collected. For example, the use of a cemented doublet, an achromatic condenser lens, or even an aspherical achromatic doublet can give higher detection efficiency compared to a long working distance objective lens system, where both the f-number and numerical aperture are sufficiently large to provide long working distances while maintaining high detection efficiency. Such relatively simple condenser lenses can be manufactured according to the state of the art (e.g., Edmund Optics, Thorlabs, Ulooptics, Newport, etc.) with much larger diameters compared to high-end objective lenses, allowing, for example, signal detection with a numerical aperture of NA = 0.53 at a working distance of several cm for aspherical doublets (singlets). Dubletts, of course, have much better control over focal position and extension over the wavelength range emitted by the fluorophores. The separation of the optics for fluorescence excitation by means of the Ring-Array illumination arrangement with a high effective synthetic aperture at very large working distances and a large variability of the positions of the individual excitation light sources, from the optics for fluorescence detection allows much higher resolution and direct integration into FIB-SEM systems and other imaging systems with specific requirements for accommodating the light-optical devices.

**Improvement of Fluorescence Collection**

In order to register the signal (e.g. fluorescence) generated in the object by the focusing or by the generation of donut (or otherwise structured) patterns by the Ring- Array illumination for imaging (e.g. confocal, STED/MINFLUX microscopy), a suitable detection system is required (see Fig. 8 for a first schematic example of fluorescence detection). In the following implementation examples, a working distance in the range of several centimeters is assumed.

In principle, for example, a lens with a numerical aperture (compare Fig. 3) would be sufficient for the detection of a fluorescence signal; however, the photon collection efficiency with such a single collecting lens is very low; for example, for NA = 0.28 it would be lower by a factor of $(1.4/0.28)^2$ ~ 25x; and for NA = 0.1 ist would be $(1.4/0.1)^2$ ~ 200x lower than with an immersion lens of aperture NA = 1.4 as typically used in SRM.

As in other laser scanning imaging systems, the resolution achievable in the Ring- Array mode depends on the local intensity distribution generated at a specific object location for fluorescence excitation; the location position is thereby determined by the position of the object stage ("stage scanning") with respect to the localized illumination (e.g. intensity maximum for focal intensity distribution, or the central zero for donut intensity). According to the state of the art, local stage positions can mechanically be determined with an accuracy σloc in the 1 nanometer range. Similar figures are obtained if the localization is performed on the basis of the focal fluorescence emission (Albrecht et al., 2002; Pertsinidis et al., 2010; Balzarotti et al., 2017; Gwosch et al., 2020). At a constant registered number of emitted photons, $\sigma_{loc}$ is inversely proportional to $(NA)^2$ (Cremer & Edelmann 2000; Cremer et al. 2017): In the object plane, this results in $\sigma_{loc}$ ~ FWHM/sqrt[ $N_{phot}$], where FWHM is the half- width of the PSF (proportional to 1/NA) and $N_{phot}$ is the number of detected photons (proportional to $NA^2$ ); hence $\sigma_{loc}$ ~ $(1/[NA)xsqrt(NA)^2]$ = $1/(NA)^2$ .

**Improvement of photon collection by dichroic mirrors**

Using the same objective lens for fluorescence excitation and detection, the localization accuracy $\sigma_{loc}$ is diminished by a factor of e.g. $(1.4/0.28)^2$ = 25x when using an objective with a large working distance (e.g., NA = 0.28) compared to an objective with a high numerical aperture (NA = 1.4) but a small working distance under otherwise identical conditions; for an objective with NA = 0.06, the localization accuracy would be even worse by a factor of $(1.4/0.06)^2$ = 544x. This is of utmost practical importance especially for fluorescence microscopic techniques such as CLSM, SMLM, MINFLUX/SIMFLUX: For example, the optical resolution achievable with single molecule microscopy techniques is proportional to the localization accuracy $\sigma_{loc}$ ; i.e. under otherwise identical conditions, if an objective with NA = 0.28 were used instead of an objective with NA = 1.4, the optical resolution due to $\sigma_{loc}$ would be a factor of 25 worse, e.g., only 250 nm instead of 10 nm; with an NA of 0.06, the resolution would be only 5.4 µm instead of 10 nm ($\lambda_{exc}$ = 488 nm).

In the following, some ways will be discussed in order to improve the efficiency of fluorescence collection when using the Ring-Array arrangement according to the invention. Generally, when using the Ring-Array arrangement in the scanning mode, the half-width (FWHM) of the focused beam in the object plane is independent of the NA of the detection optics (Fig. 3, 8) and corresponds, for example, to an effective numerical aperture $NA_{array}$ ~ 0.94 for the assumptions made in Table 1 ($\lambda_{exc}$ = 488 nm; n= 1; $\alpha_{min}$ = 45°; $\alpha_{max}$ = 70°). In this case, in principle the optical resolution would not depend on a high photon yield emitted from the object but mainly on the positioning of the Array-Beam (focused/donut) and its PSF (related to $NA_{array}$); however, also in this case, the photon yield registered from the individual sites excited by the scanning Ring-Array illumination beam is substantial for contrast and hence for effective resolution (Stelzer, 1998). Therefore, for a full use of the Ring-Array microscopy concept, it remains important to enhance the fluorescence collection as much as possible. In the SMLM mode, the resolution depends essentially on the localization precision directly correlated to the photon yield.

In the following, some possibilities of fluorescence collection are outlined in order to improve the efficiency of fluorescence detection when using the Ring-Array arrangement. A general schematic example is given in Fig. 8: Here, photons emitted from the object plane at a very large working distance are collected with the help of suitable dichroitic mirrors with maximum transmission for the excitation wavelength(s) and maximum reflection for the fluorescence emission of the object as well as with the help of one or more detectors (e.g. "point detectors" or area sensors (CCD, sCMOS). This scheme is only intended to convey that an efficient photon collection is possible also at a very large WD. Many other arrangements are also feasible.

**Improvement of photon collection by low numerical aperture lenses**

Instead of using a single objective lens with low numerical aperture to detect the local fluorescence emission generated by the Ring-Array arrangement according to the invention in the focusing mode, an implementation according to this concept consists of an array of multiple objective lenses with low NA arranged at a correspondingly large working distance around the ring array (Fig. S5). The individual fluorescence signals registered by these with the aid of downstream detectors are added up to a total signal (total number of detected fluorescence photons) and used for imaging in the scanning mode (focused/donut). As mentioned, this is possible because the positional information essential for image resolution is given by the location **O** of the Ring-Array generated focus/donut intensity distribution. For example, if a single detection lens has an NA of

0.25, this results in an aperture angle of $\alpha$ = 14.5° in vacuum, corresponding to a solid angle of $\Omega = 2\pi(1 - \cos \alpha) = 0.06\pi$. To achieve the same photon collection efficiency as a lens with $NA_{det}$ = 0.92 (vacuum) ($\alpha$ = 67.5°; $\Omega$ = 1.23π, about 10 of these collection lenses would thus have to be arranged at a working distance of e.g. 3.4 cm. Such arrangements appear to be implementable in principle: Since fluorescence emission is largely isotropic, this may be done also in the space marked by the solid angle $\Omega_{bottom}$ (Fig. 2). For example, assuming a ring conformation with a radius of 5 cm, a section of 31cm/10 = 3.1 cm is available on a circle of 2π x 5 cm with a circumference of 31 cm per collecting lens. Compared to the collection efficiency of an immersion objective with NA = 1.4, the total fluorescence signal (proportional to the number $N_{phot}$ of all registered fluorescence photons) in this case is only reduced by a factor of $(1.4/0.9)^2 = 2.4$, a loss in collection efficiency that can be compensated, for example, by longer acquisition times. When using the single-molecule mode according to the Ring-Array arrangement (assuming coherent detection), the localization precision $\sigma_{loc}$ is reduced by an overall factor of about 2.4 compared to the prior art (immersion lens with NA = 1.4) under the assumptions made.

**Improvement of photon collection by optical fiber arrays**

Instead of a single objective lens with a large working distance and correspondingly low NA for signal detection, also an array of endoscopic optical fiber arrays may be used. Since the spatial distribution of the light sources in the Ring-Array of the invention is largely variable without adversely altering the intensity distribution required for high resolution, the optical fiber arrays can be implemented in a variety of ways. For example, the optical fibers may be implemented in a number of bundles without disturbing the Ring-Array arrangement or the devices required for FIB-SEM or other correlative microscopy applications. As in the previous example, the position of the illuminated object is given by the adjustment (position, direction, polarization, etc.) of the scanning beam, while the optical resolution (discrimination of two fluorescence point sources) depends on the focal diameter of the scanning beam or, more generally, on the intensity distribution at the observation location.

If we assume an aperture cross-section of $(5\ mm)^2$ at a large working distance of e.g. 3.4 cm for a single glass fiber bundle as an example, this results in a solid angle (referred to the focus center **O**) of $0.5 \times 10^{-2}\ \pi$ for the aperture. Accordingly, to achieve a collection efficiency of a lens with the numerical aperture (vacuum/air) of 0.92 ($\Omega$ = 1.23π), about 250 individual bundles of optical fibers have to be positioned on a ring at a distance of, e.g., 3.4 cm instead of the lenses. The fluorescence signals recorded by the individual glass fiber bundles can then be transferred to a single detector.

**Improvement of photon collection by sensor arrays**

As mentioned above, there are many ways to distribute the sources Si in the Ring-Array (Fig. 4). One of them is locate the sources of different „array rings“ in concentric rings, separated by material transparent to the fluorescence emission of the object (Fig. S5a). These transparent concentric rings can then be used for collection of fluorescence photons.

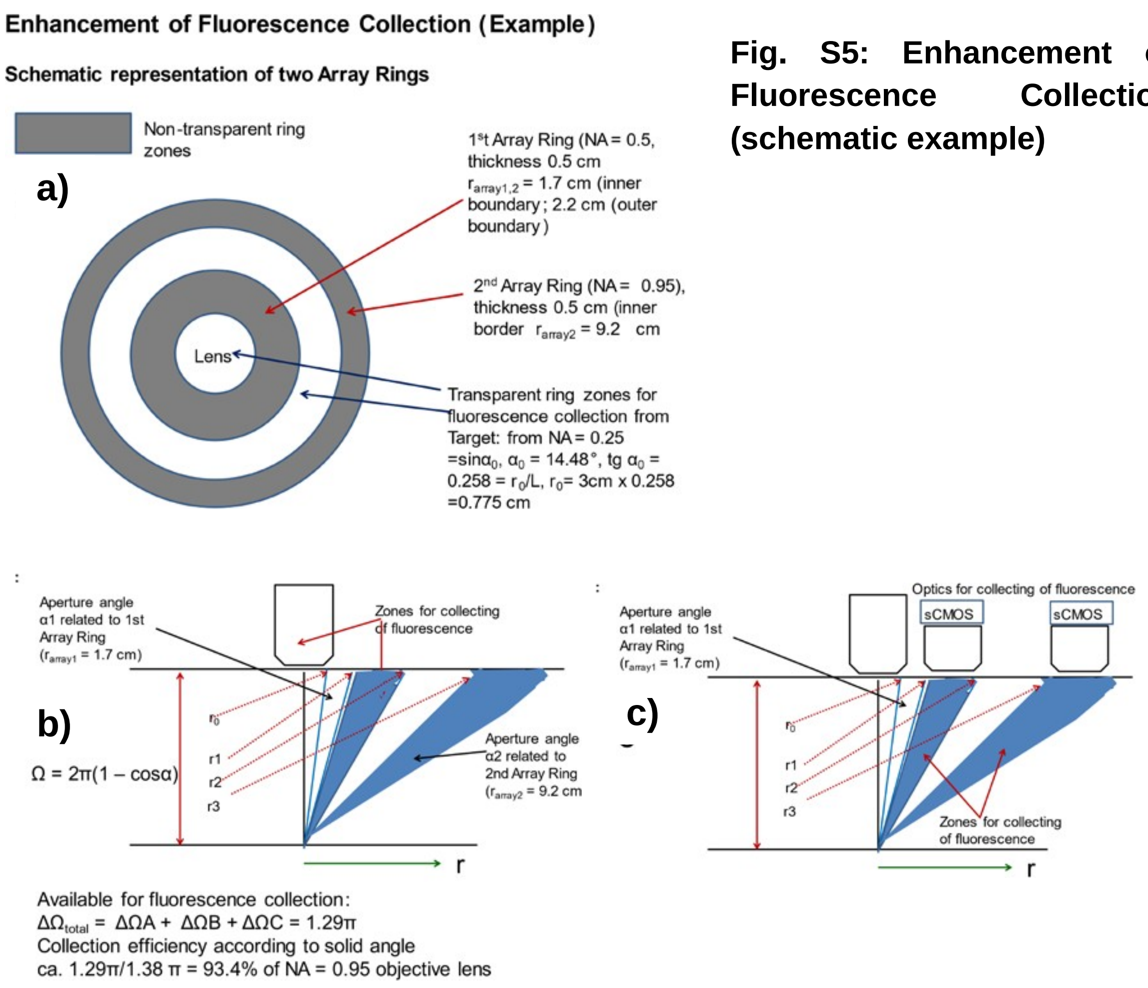


**Fig. S5: Enhancement of Fluorescence Collection (schematic example)**

**Fig. S5: Enhancement of Fluorescence Collection Efficiency (schematic example).**

*In this example for the improvement of fluorescence collection efficiency, let us assume (Fig. S5a) an Array-Ring with an the inner array-ring corresponding to a synthetic aperture NA ~ 0.5, and an outer array ring corresponding to a synthetic aperture NA ~ 0.95 ($\alpha_{max}$ ~ 70°). The zones between the inner and the outer array-ring are transparent for fluorescence excited in the object by the Ring-Array beams (Fig. S5b). Closely above these transparent ring zones, the fluorescence emitted is collected by an array of microlenses which guide the light to appropriate detectors, e.g. sCMOS sensors (Fig. S5c); the sum of all the sensor signal is then a measure for the fluorescence yield at the object site identified by the scanning Ring-Array illumination beam.*

A still more radical solution would be to place a large number of sCMOS sensors directly above the transparent ring zones (without lenses); the signals registered would be computationally summarized and serve as a measure for the fluorescence emission of the object at the scanning site. For example, assuming a Ring-Array collecting efficiency corresponding to a Numerical Aperture around 0.9, an example estimate gives a total transparent ring zone area of about 7,000 $mm^2$ . At first sight, this appears to be far beyond economic realization; however, using economic sCMOS sensors for Smartphones sufficient for single molecule localization microscopy (Diederich et al. 2020), such an option would cost not more than a high performance scientific CCD camera some years ago. The advantage of such a solution would be its simplicity. In combination with appropriately bright fluorophores, for example nanographenes (Liu et al., 2020), such a fluorescence collection system would in principle be sufficient for various microscopy modes (focused, STED, SIM) established for high NA/low WD microscopy; in combination with coherent fluorescence collection modes (Hell et al., 1994b), also the implementation of SMLM, MINFLUX and SIMFLUX modes should be feasible.

It may be noted that in the case of Ring-Array based nanosizing using a low NA/large WD objective lens for fluorescence detection (Fig. S3) and appropriate labeling conditions, in many cases the photon collection efficiency of the low NA lens should already be sufficient. For example, assuming a low NA = 0.28 with WD = 3.4 cm, the fluorescence collection efficiency of such a lens compared to a NA = 1.4 lens (WD = 0.17 mm) is smaller by a factor $(1.4/0.28)^2$ = 25. This means that instead of e.g. 5,000 photons registered from a single fluorescent molecule at NA = 1.4 (Liu et al., 2020), only 250 would be detected at NA = 0.28. Numerical simulations of SMI nanosizing with respect to the photon efficiency required (Cremer et al., 2003) indicate, however, that already a total of ~ 1,000 photons registered from a nanostructure should allow a correct nanosizing down to the 100 nm range. Hence, labeling the nanostructure with a few fluorophores should be sufficient.

**Scanning Options**

In the examples described for the practical implementation of Ring-Array microscopy, the scanning of the specimen at selected sites for SRM imaging was assumed to be performed by point-by-point scanning. Since the regions-of-interest of such selected sites to be scanned are typically very small (in the micrometer range), scanning by moving the specimen ("stage scanning") can be extraordinarily fast.

However, by suitable adjustment of the illumination light sources, it might also be possible to perform beam scanning, thus greatly accelerating the imaging process. This would be realized by coordinated changes in the directions of the collimated beams while maintaining their phase relationships. To minimize the adjustment effort, the light sources in the ring array may be arranged in a number of "subarrays" that are closely interconnected.

When applying the present concept to larger objects with spatially widely distributed fluorescent emitters (e.g., a cellular spheroid containing GFP (green fluorescent protein)-labeled histones or immunolabeled receptors), it is important to consider that the multiple beams emitted by the Ring-Array will also excite fluorophores outside the focal region; this emission may become so strong that the fluorescence generated in the focal region may not be clearly distinguished from the fluorescence of this "background." One way to overcome this problem is to use two-photon excitation. Such an implementation, however, requires additional adjustment effort due to the limited coherence lengths of short laser pulses.

In principle, the use of coherent light sources allows the implementation of interferometric signal detection as well: the generation of synthetic apertures can enable the extraction of medium and high resolution images using low NA microscope lenses (Mico et al. 2006). In contrast to the Ring-Array illumination concept using coherent light sources, incoherent illumination of semi-transparent samples has been shown to enable quantitative phase recovery using quadri-wave lateral shear interferometry (Bon et al. 2014). In combination with a range of polarization-sensitive detectors (e.g. by adding appropriate polarization filters, a Ring-Array illumination arrangement may also be used to multiplex ellipsometric measurements (Egner 2002).

In the original concept of a confocal laser scanning 4π-fluorescence microscope providing "super-resolution" at a large working distance (Cremer and Cremer 1978), it was assumed that point-by-point excitation of the object in three dimensions (3D) was possible by using "4π-holograms" to focus the incident coherent light beams onto a focal "spot" with a diameter smaller than that obtainable by focusing through an objective lens with the same working distance. In the Ring-Array arrangement (Fig. 1-3,8), a suitably high number N (up to several hundred) of coherent, continuously emitting point sources $S_1$ , S2 ,..., $S_N$ is assumed. The point sources $S_1$ , . ,$S_N$ have here fixed phase and polarization relations to each other. For an arbitrary but fixed configuration (positions, phases, polarizations, propagation directions, intensities) of light sources, the origin of the coordinate system is placed on the theoretical absolute maximum of the focal illumination intensity distribution (i.e., the theoretical "focus" **O**, see Fig. 1). The origin point **O** (x,y,z = 0), together with the center of gravity of the positions of the light sources in the Ring-Array arrangement defines a set of two positions in space. The line through both points forms the optical axis (z). In contrast to focusing by 4π- holograms (Cremer&Cremer 1978), according to the microscopy concept presented here, it is possible to regulate the direction, intensity, polarization and phase for each of the coherent illumination beams emitted by a Ring-Array arrangement individually and independently of the others.

**STED/MINFLUX imaging with the Ring-Array arrangement**

As another embodiment of the present Ring-Array microscopy concept, the ring array arrangement enables the generation of a STED depletion beam with an adjustable donut mode or a donut pattern that can be used in MINFLUX mode. Figs. 6c-e, Table 2 Maintext, and Fig. S5 show that such donut-shaped intensity distributions can be realized with suitable Ring-Array arrangements with parameters corresponding to a high numerical aperture objective according to the state of the art (Hell et al. 1999; Hell 2009; Sahl & Hell 2019). The image acquisition is performed by object scanning with the Ring-Array generated intensity distribution.

a)

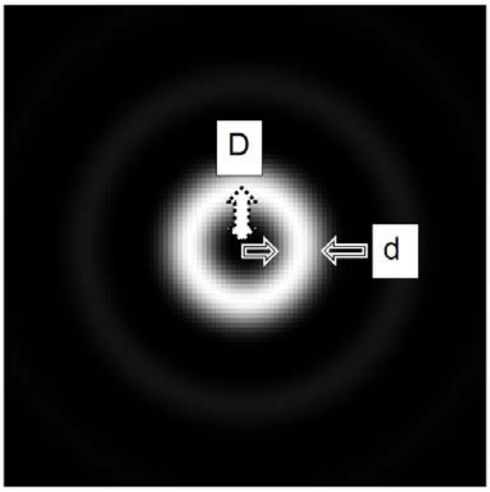


b)

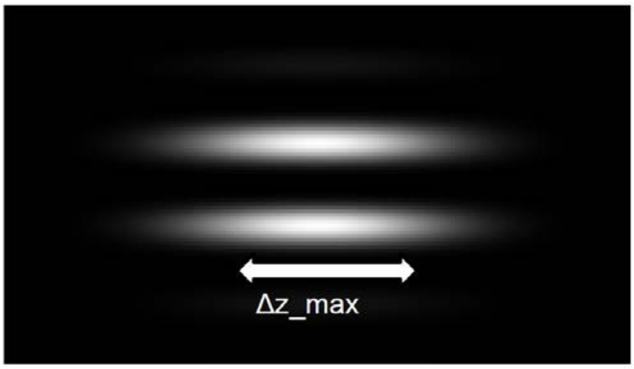


**Fig. S6: Characteristic dimensions of a Ring-Array illumination generated donut intensity distribution**

*D: Width of the central lateral "gap" (from the central minimum to the maximum of the intensity of the torus ring ("bulge") in the object plane (xy);*

*d: Half-width (FWHM) of the donut ring intensity;*

*Δz_max: half-width (FWHM) in the direction (z) of the optical axis at maximum intensity.*

*When Ring-Array illumination is used, for D values around 0.34 $\lambda_{exc}$ (ice, vacuum/air) are calculated; these figures are very similar to those obtained when using an objective lens with a large numerical aperture (half aperture angle α = 70°, NA = 0.94 (n=1 (Table 2).*

*For Δz_max, values around 2.5 $\lambda_{exc}$ (or $\lambda_{STED}$, $\lambda_{MINFLUX}$) are obtained in vacuum/air, or in ice. These estimates are larger by about one wavelength unit than when using an objective lens with NA = 0.94 (Table 2).*

*For an angle of e.g., $\alpha_{max}$ = 70° that can be realized according to the Ring-Array concept (see Fig. 1) this angle corresponds to a numerical aperture (NA) of an objective of 0.94 (vacuum, n= 1), as used e.g. in STED/MINFLUX microscopy according for the generation of ("donut") intensity distributions; in contrast to such lens-based objectives with only small working distance (typically in the range of 0.2 mm), in Ring-Array microscopy the working distance (WD) can be increased up to* *the range of several cm (e.g. WD = 3.4 cm, or 5 cm), while maintaining almost the same resolution.*

The Ring-Array microscopy concept makes it possible to generate a specific local intensity distribution (focusing, donut) at a specific location (xyz) in the region of the object plane even at extremely long working distances, to excite the fluorescence of a selected ROI, and to image this with the aid of suitable detector devices (see Fig. 9 for a schematic representation of a detector device).

To generate an image of the ROI selected using such local intensity distributions, the ROI is scanned point-by-point using state-of-the-art laser scanning microscopy methods (Cremer & Cremer 1978; Sheppard & Wilson, 1978; Brakenhoff et al., 1979, 1985; Masters, 1996), and the associated fluorescence signal is measured point-by- point. According to the state of the art, the scanning can be performed either by shifting the object coordinates ("stage scanning") or by moving the focused beam ("beam scanning"). However, there are some specific requirements to be considered when using such scanning methods in Ring-Array microscopy, for example in FIB- SEM:

In preparation for the analysis of cryosamples in the FIB-SEM, it is useful to first scan a sample of e.g. 1 mm diameter with an objective lens of relatively low resolution in order to delimit the regions of interest. Such a region of ~ 1 $mm^2$ would then suffice for further analysis by high resolution. The target region isolated for this purpose (e.g. a lamella) should be e.g. approx. 500 nm thick and approx. 20 to 50 µm wide, whereby a light-optical resolution of 50 - 100 nm would be desirable.

This results in various requirements for the screening:

Coarse scanning of an object area of approx. 1,000 µm diameter: Let us assume that an object resolution of e.g. 1 µm is desired for this; the Numerical Aperture required (e.g. NA ~ 0.3) may be obtained either by a glass lens (if the sterical conditions allow), or also by Ring-Array illumination with an array-ring distribution of sources (Fig. 4) corresponding to such a low NA. An object with an area of $(1{,}000\ \mu m)^2$ is to be scanned in steps of 0.5 µm, i.e. a total of $(2{,}000)^2$ steps of 0.5 µm step size each. Using sensitive detectors with a dwell time per step of 0.1 ms, the scanning time would be (2,000)2 s = 400 s (about 7 min); the speed of the object stage (scanning in total a line of 1,000 x [1,000/0.5] µm = 2,000 mm length) during "stage scanning" would be vstage ~ 5 mm/s, and hence feasible according to the state of the art (e.g. https://www.thorlabs.com/newgrouppage9.cfm?objectgroup_id=5360).

These example estimates show that even such large object areas can be scanned with a practicable expenditure of time.

For the much smaller step sizes required for the subsequent SRM scanning at enhanced resolution of selected ROIs with typical dimensions in the < 100 µm range, a second scanning stage device with nanometer resolution may be applied. If the object area (e.g. 1 $mm^2$) is limited by the coarse screening described above to an object area of e.g. $(20\ \mu m)^2$, and a fine screening is now to be performed at an enhanced resolution (MINFLUX/STED) of e.g. 20 nm, a step size of 10 nm is required; such step sizes can be achieved without difficulty with piezo stages according to the state of the art (e.g. Hii et al., 2010; https://www.physikinstrumente.de/)

Comparable estimates of the applicability of the Ring-Array arrangement for high-resolution scanning microscopy with extremely large working distance are also valid for serial block-face SEM (room temperature in epoxy or metacrylate resin), and are also applicable to other microscopy techniques with large working distance, such as direct Correlated Lightoptical Microscopy (dCOLM). These include, for example, applications in biomedical engineering, such as high-resolution light-optical analysis of three-dimensional cell assemblies (organoids,spheroids); in contrast to [Birk et al., 2017], the Ring-Array arrangement according to the present concept would open up the possibility of using the free solid space $\Omega_{cente}$r (e.g., 0° ≤ α ≤ 45°) according to Figs. 1,2,8 to devices for manipulation, with otherwise almost the same resolution. Similar applications could arise in materials analysis/machining.
